# Martini 3 coarse-grained models of azobenzene-based photolipids: Modulation of membranes with light

**DOI:** 10.64898/2025.12.05.692541

**Authors:** Cristina Gil Herrero, Thilo Duve, Sebastian Thallmair

## Abstract

Photoswitchable lipids offer an attractive way to manipulate the biophysical properties of membranes by means of light. Their application to modulate membrane properties, manipulate membrane proteins, and photocontrol cargo release is gaining popularity. Here, we present coarse-grained Martini 3 models for azobenzene and azobenzene-based photoswitchable lipids. Our models show good agreement with atomistic reference simulations. Furthermore, we apply our coarse-grained photolipid models to study photocontrol of lateral phase separation, protein flexibility, and membrane permeability. The results agree well with experimental data from the literature and highlight the broad applicability of our Martini 3 photolipid models. They will enable studying the impact of photolipid switching on large membrane systems as well as on their (bio)molecular interaction partners.

## 1 Introduction

Lipid membranes play a fundamental role in biology, forming barriers and serving as the border of cells, and in this way separating the cells’ interior from their external environment. Also, many organelles such as the endoplasmic reticulum or mitochondria, are surrounded by a membrane. Based thereon, membranes are crucial for the uptake of molecules from the outside as well as in cell-cell communication, for instance at synapses, where the formation and fusion of lipid vesicles is involved. But lipid membranes do not only serve as a border, they also constitute the natural environment of membrane proteins. They are able to regulate transmembrane proteins, for instance, via their thickness or tension.^1,2^ In addition, individual lipids may serve as specific protein ligands.^3–6^ While natural lipids exist in a tremendous variety, changing their biophysical properties requires chemical reactions such as the reduction of C=C double bonds in a lipid tail, which would result in a more ordered, saturated lipid.

Artificial photolipids, which contain a photoswitchable moiety in at least one of their tails, allow manipulation of membrane properties on a very fast time scale by means of light without physical intervention.^7–10^ A commonly used photoswitch in many biological applications is azobenzene.^11^ Azobenzene has also been incorporated into lipid tails, resulting in azobenzene-based photoswitchable lipids.^10^ In recent years, these photolipids have been applied to study the modulation of biophysical properties of membranes as well as their impact on proteins embedded in such membranes. It has been shown that photoswitching azoPC, the prototypical photoswitchable lipid, from *trans* to *cis* resulted in an increase of the area per lipid, membrane thinning and vesicle instabilities.^12,13^ Furthermore, a ternary lipid mixture containing azoPC allowed photocontrol of lateral phase separation in giant unilamellar vesicles.^14^ In earlier studies, phosphatidylcholine (PC) lipids with two photoswitchable tails were used to photocontrol cargo release from liposomes, focusing on potential applications for drug delivery and controlled release.^15,16^ More recently, also azoPC-containing liposomes have been investigated for their use in light-controlled release.^17^ The impact of azoPC photo-switching on membrane order and dynamics has been investigated in detail using solid-state NMR measurements.^18^ In the presence of the transmembrane protein diacylglycerol kinase (DgkA), this impact was further increased, and in addition, changes in the protein dynamics were observed. In globotriaosylceramide (Gb3) glycosphingolipids, it was shown that the position at which azobenzene moieties are incorporated into the lipid tails determines how photoswitching influences lateral phase separation.^19^ Moreover, the peripheral membrane protein Shiga toxin, which binds to the headgroup of Gb3 lipids, showed changes in membrane binding depending on the switch state of the photoswitchable Gb3 lipids.^19,20^ Using diacylglycerol lipids with two photoswitchable tails allowed to photocontrol the opening probability of the mechanosensitive ion channel gramicidin A.^21^ Switching from *trans* to *cis* increased the conductance of gramicidin A by more than 50%. These examples highlight the diverse applications of photoswitchable lipids and their potential for drug delivery and as chemical probes for membrane biophysics.

Molecular dynamics (MD) simulations are an important technique to characterize biomolecular ensembles, providing complementary information to experiments. Nowadays, they are a key tool in molecular biology, offering access to unprecedented structural and dynamic information of biomolecular systems. While the light-induced switching of photolipids involves electronically excited states and thus, an accurate treatment of the ultrafast switching requires quantum mechanical treatment, the modeling of the two (meta)stable states can be performed in the electronic ground state using commonly available classical force fields. A detailed parametrization of photoswitchable fatty acids was performed for the atomistic CHARMM36 force field.^22^ Based thereon, photolipid parameters were generated to study membrane properties and membrane permeability in the two switch states of azoPC.^23,24^ Similarly, CGenFF-based azobenzene parameters were combined with CHARMM36 lipid parameters to study photolipid properties and membrane permeability.^25^ In other studies, atomistic MD simulations of membrane patches containing azobenzene-based photolipids were performed using the General Amber Force Field (GAFF).^13,26^

While atomistic MD simulations provide detailed structural information of photoswitchable membranes, large-scale simulations, for instance of phase separation, are still computationally challenging. Here, coarse-grained (CG) models such as the Martini force field,^27^ have been proven highly valuable for achieving the required sampling at a reasonable computational cost. However, while CG models of two molecular motors and the molecular switch oxindole were presented recently,^28^ CG Martini 3 models of photoswitchable lipids are not available to the best of our knowledge. Here, we present the CG Martini 3 models of azobenzene as well as azobenzene-based photolipids and fatty acids. Their parametrization is based on the CHARMM36 models of photoswitchable fatty acids.^22^ We incorporated the azobenzene moiety into the CHARMM36 lipid parameters of 1,2-distearoyl-sn-glycero-3-phosphatidylcholine (DSPC) and the resulting lipid models served as a reference for our CG models of PC lipids with one and two photoswitchable tails.

The paper is structured as follows: After the Computational Methods section, we first present the CG azobenzene model and its parametrization. This is followed by the CG models for the photolipids as well as the photoswitchable fatty acids. We then demonstrate the applicability of our CG models in several examples, including studying lateral phase separation in binary and ternary lipid mixtures, the impact of photoswitching on the transmembrane protein DgkA, and the membrane permeability of the drugs salbutamol and baricitinib. Before concluding, we discuss the limitations of the presented CG models.

## 2 Computational Methods

All simulations containing biomolecular ensembles, i.e. lipid membranes and optional proteins, were performed with the program package GROMACS (version 2020.4 and 2024.5).^29,30^ The only simulations performed with a different software are the reference simulations for *trans-* and *cis-*azobenzene, which were performed with Bartender based on xtb.^31,32^

### 2.1 Semi-empirical quantum chemical simulations

Semi-empirical quantum chemical simulations were used as reference for the parametrization of the azobenzene CG models. The starting structures used for these calculations were obtained using the all-atom (AA) force field CHARMM36m. The azobenzene topologies were obtained from CHARMM-GUI’s Ligand Reader & Modeler based on the CHARMM General Force Field (CGenFF).^33–35^ Azobenzene was minimized in vacuum while constraining the switching dihedral (ϕ*_C_*_−_*_N_*_=_*_N_*_−_*_C_*) to -15° and 180° for the *cis* and *trans* isomers, respectively. The minimized structures were then used to run 100 ns simulations using the GFN2-xTB method with an implicit solvent model for water^36,37^ at the default temperature of 310.00 K to allow for more efficient sampling of the conformational space of the molecule.

### 2.2 All-atom simulations

The AA simulations were performed with the CHARMM36 force field^38^ and were employed as reference for the parametrization of the CG models. The CHARMM36 models of the photoswitchable lipids were built by combining the optimized azobenzene CHARMM parameters for a protonated fatty acid of Klaja et al.^22^ with the default lipid and deprotonated fatty acid parameters in CHARMM36. Atomistic membrane systems were constructed by embedding the photoswitchable lipids into a pre-equilibrated DSPC:1-palmitoyl-2-oleoyl-sn-glycero-3-phosphocholine (POPC) (1:3) membrane patch of 8 lipids in total. The photoswitchable lipids were aligned to the DSPC lipids in the upper and lower leaflet using Visual Molecular Dynamics (VMD)^39^ and inserted into the patch by replacing the DSPC coordinates. The resulting bilayer, containing one photoswitchable lipid and three POPC lipids per leaflet, was replicated 4×4 in the xy-plane to reach the desired system size (32 photoswitchable lipids and 96 POPC lipids). The systems were solvated with 2750-2950 water molecules using the CHARMM-modified TIP3P water model, neutralized, and 0.15 M NaCl was added (37/69 Na^+^ ions for the neutral/charged systems, respectively, and 37 Cl^−^ ions). The final system size was approximately 7 × 7 × 8.5 nm^3^.

We applied the recommended simulation settings taken from CHARMM-GUI^40–42^ for the CHARMM force field. The minimization was performed for 5000 steps using the steepest descent algorithm, and the equilibration was divided into six steps to gradually release the position restraints in the membrane and to increase the timestep from Δt = 1 fs to Δt = 2 fs. The first three equilibration steps were carried out in an NVT ensemble for 125 ps each, and the last three in an NPT ensemble for 500 ps each. The temperature was maintained by applying a Berendsen thermostat^43^ (τ*_T_* = 1 ps); the pressure by applying a Berendsen barostat^43^ (τ*_p_* = 5 ps). We performed the AA reference simulations at two temperatures, namely T_ref_ = 303.15 K and T_ref_ = 330.15 K, and a pressure of p_ref_ = 1 bar. For the production run of 1 µs, a timestep of Δt = 2 fs was employed, and the temperature and pressure were maintained with a Nośe-Hoover thermostat^44^ (τ*_T_* = 1 ps) and a Parrinello-Rahman barostat^45^ (τ*_p_* = 5 ps), respectively. Van der Waals interactions were treated with a cutoff scheme (r_c_ = 1.2 nm); Coulomb interactions were calculated using Particle Mesh Ewald (PME) (r_c_ = 1.2 nm).

### 2.3 Coarse-grained simulations

All CG simulations were performed using the Martini 3 force field (version 3.0.0).^27^ The CG models of the photoswitchable lipids were built by combining the optimized azobenzene parameters for both *trans-* and *cis-*isomers with the refined Martini lipid building blocks (version 3.1),^46^ as described in Section 3.2.

For the CG simulations of the azobenzene photoswitch, the molecule was placed in a simulation box with an edge length of 4.5 nm and solvated in water using *gmx solvate*. To build the CG membrane systems, we used the CG system builder COBY.^47^ The CG membranes contained the same amount of lipids as the AA reference membranes, namely 32 photoswitchable lipids and 96 POPC lipids (ratio 1:3). The membranes were solvated with approximately 1500 water beads, 0.15 M NaCl was added (approximately 18/50 Na^+^ ions for the neutral/charged systems and 18 Cl^−^ ions), and additional Na^+^ ions were added when necessary to neutralize the systems with deprotonated fatty acids. The final system size was approximately 6.2 × 6.2 × 8.5 nm^3^.

For all Martini CG simulations, an initial steepest descent minimization of 2000 steps was performed, followed by a single equilibration in the NPT ensemble employing the v-rescale thermostat^48^ (τ*_T_* = 1.0 ps), and the Berendsen barostat^43^ (τ*_p_* = 5 ps). A timestep of Δt = 10 fs was used during the equilibration for a total equilibration time of 500 ps. Van der Waals and Coulomb interactions were treated with a cutoff scheme (r_c_ = 1.1 nm) and the Potential-shift-Verlet was employed as Coulomb modifier. Most of the simulation parameters were kept for the production runs, except for the barostat, which was changed to a Parrinello-Rahman barostat^45^ (τ*_p_* = 12 ps). Also, the timestep was increased to Δt = 20 fs for the 1 µs long production runs. Like the AA reference simulations, the CG simulations were performed at two temperatures, namely T_ref_ = 303.15 K and T_ref_ = 330.15 K, and a pressure of p_ref_ = 1 bar. Simulations of azobenzene were performed at T_ref_ = 310.00 K in accordance with the semi-empirical reference simulations.

### 2.4 Analysis

To perform the analysis, all trajectories were initially processed with the GROMACS tool *gmx trjconv -pbc whole* to avoid molecules being split across the borders of the box. The bonded terms of the pseudo-CG (mapped from AA) and CG simulations, namely distances, angles, and dihedral angles, were calculated using the MDAnalysis library,^49^ ^50^ based on the center-of-geometry mapping as recommended for Martini 3.^27^ For angles and dihedral angles, the functions *calc angles* and *calc dihedrals* were used, respectively. The bonded terms were evaluated for each photolipid.

Analysis of the solvent accessible surface area (SASA) was conducted using *gmx sasa*. The default parameters recommended for Martini 3 were used, with 4800 grid points (*-ndots*) and a probe radius of 0.191 nm (*-probe*), corresponding to the van der Waals (vdW) radius of a tiny bead.^51^ The AA trajectories used vdW radii described by Rowland and Taylor,^52^ while CG trajectories used 0.191 nm, 0.230 nm, and 0.264 nm for tiny, small, and regular beads, respectively.

The octanol-water partitioning was estimated from the solvation free energies of the respective molecule in both solvents, which were calculated by performing a thermodynamic integration (TI) using GROMACS (for details, see Section 1 in the Supporting Information). The TI was analyzed using the Python package Alchemlyb.^53^

To compute the angle between the azobenzene moiety and the membrane normal, first, planes were fitted to the PO4 (CG) or P (AA) particles in each leaflet. Points above and below a defined z-split (around z = 0 nm) were used to generate the upper and lower leaflets. For each leaflet, the phosphate atom positions were used to perform a least-squares fit to a plane of the form z = ax + by + c. The normal vector of each plane was then computed analytically from the plane coefficients as n⃗ = [−a, −b, 1]. The average of these two normals defined the overall membrane normal vector. The angle was then calculated between the membrane normal vector and the vector connecting the two azobenzene rings. For the AA model, this vector was defined between the centers of geometry of all carbon and hydrogen atoms in each aromatic ring. For CG, the corresponding vector connected two virtual sites (V8 and V9) placed at the centers of the rings. All properties described above were computed for each frame of the trajectory, and histograms were generated to compare the photolipids’ behavior at the different resolutions.

The GROMACS tool *gmx density* was employed to compute the mass densities along the z-axis, with selections including the whole membrane (POPC and the photolipids), the PO4 beads, and the photolipids only. For visualization, the photolipid densities are scaled by a factor of two in the plots.

The program FATSLIM^54^ was employed to compute the area per lipid using the P atoms at AA and the PO4 beads at CG resolution for the phospholipids; for the fatty acids, the COO^−^/COOH bead, or the carbon atom of the carboxylate/carboxyl group was used. For the computations of the membrane thickness, first, the indices of the P atoms/PO4 beads were extracted and split into the upper and lower leaflet. The z-component of the distance between the center of mass (COM) of both groups was then obtained using the GROMACS tool *gmx distance*, and the average and standard errors were calculated using *gmx analyze*.

### 2.5 Application setups

#### 2.5.1 Azobenzene interaction with POPC membrane

To study the interactions between the isomers of azobenzene and a membrane, a POPC bilayer was solvated in water and 0.15 M NaCl. 30 azobenzene molecules were placed randomly in the solvent phase using COBY’s flood feature.^47^ The final system size was approximately 7.9 × 7.9 × 16.5 nm^3^. One replica of 600 ns was run per isomer, while the first 100 ns were considered as further equilibration. The remaining 500 ns were used for density calculations.

#### 2.5.2 Binary and ternary lipid mixture

To investigate lipid interactions and experimentally observed light-controlled membrane domain formation, we performed CG simulations of bilayers composed of 1,2-diphytanoyl-sn-glycero-3-phosphocholine (DPhPC)^55^, 1-stearoyl-2-(p-butyl-p-botanoyl-*trans-*azobenzene)-phosphatidylcholine STAPC, and cholesterol^56^ at a 4:4:2 ratio. Initial systems were constructed with 100 lipids per leaflet (including cholesterol). All simulations were conducted at 303 K. After energy minimization (2000 steps, steepest descent) and a 1 ns equilibration, a 1 µs production run was carried out to equilibrate the initial patch. This bilayer was then replicated three times in x and y direction using the GROMACS tool *gmx genconf*, resulting in a final system containing 1800 lipids in total and a box size of approximately 23 × 23 × 16 nm^3^. The larger system underwent another minimization and 1 ns equilibration (Δt = 10 fs), followed by a 5 µs production simulation (Δt = 20 fs). To model the effect of *trans-cis* photoisomerization, the force field parameters were switched from STAPC to SCAPC by replacing the .itp files at the end of the 5 µs simulation. A short 50 ps long equilibration using a timestep of Δt = 1 fs was performed following minimization to ensure a smooth transition, after which the production simulation was performed for 5 µs (Δt = 20 fs).

Furthermore, we prepared a binary lipid mixture composed of 1,2-di-(p-butyl-p-botanoyl-*trans-*/*cis-*azobenzene)-phosphatidylcholine (D(T/C)APC) and POPC at a 1:3 ratio. The bilayer was built using COBY,^47^ resulting in a final system containing 1924 lipids and a box size of approximately 24 × 24 × 17 nm^3^. All simulations were conducted at a temperature of 330 K. The energy minimization was performed for 2000 steps, followed by an equilibration of 500 ns (Δt = 10 fs). All other simulation settings were identical to those used for the ternary lipid mixture.

Analyzing domain formation. Domain formation was assessed by analyzing contacts between PO4 beads of STAPC/SCAPC and DPhPC, and DTAPC/DCAPC and POPC, as well as among photolipid PO4 beads using the GROMACS tool *gmx mindist*. While the analysis was performed for the whole membrane at once, the use of the PO4 beads together with a cutoff of 0.7 nm inherently distinguished between the upper and lower leaflet. The relative contact fraction was used to quantify the degree of clustering:^57^

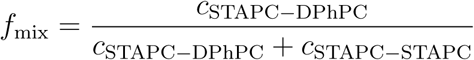

where *c* represents the number of contacts between the two lipid species indicated in the subscript.

#### 2.5.3 Transmembrane protein DgkA embedded in photoswitchable membrane

To study the influence of photolipid switching on transmembrane proteins, we simulated the transmembrane protein DgkA in a photoswitchable membrane. The crystal structure of the DgkA trimer (PDB ID: 4UXX) was preprocessed by removing non-protein molecules and the connectivity records. The individual protein chains were separated, and the most complete chain (chain B) was used to generate a GōMartini 3 protein model^58^ with Martinize2.^59–61^ The secondary structure was assigned using DSSP,^62^ and virtual interaction sites were introduced at the BB beads for the GōMartini 3 model. The dissociation energy of the Lennard-Jones potentials of the GōMartini 3 model was set to ɛ_LJ_ = 9.4 kJ/mol, the default value for the GōMartini 3 model. To generate the CG structure of the DgkA trimer, the CG structure of chain B was fitted to the other two chains in the atomistic structure of the trimer. Subsequently, the CG trimer was embedded in a pre-equilibrated lipid bilayer with a composition of POPE:POPG:PTAPC = 24:6:5 in both leaflets using COBY.^47^ The membrane composition was chosen based on experimental reference data.^18^ The total number of lipids was 692. The resulting membrane was solvated with approximately 20,000 water beads, neutralized, and 0.15 M NaCl was added (352 Na^+^ ions; 227 Cl^−^ ions). The final system size was approximately 15 × 15 × 15 nm^3^. The system was minimized (2000 steps) and equilibrated for 0.5 ns (Δt = 10 fs), followed by a 10 µs production simulation (Δt = 20 fs). Three replicates were simulated at 303 K.

Analyzing protein flexibility. To compare the flexibility of the DgkA trimer embedded in membranes containing the photolipids in either *trans-* or *cis-*conformation, we calculated the root-mean-square fluctuations (RMSF) of the backbone beads using an iterative alignment protocol. The first 1 µs of each trajectory was considered as further equilibration and excluded from the analysis. The remaining trajectory from 1–10 µs was divided into 50 ns windows, each of which was processed separately to ensure the protein remained whole, centered, and aligned in the simulation box. The backbone RMSF was calculated with the GROMACS tool *gmx rmsf* after iteratively fitting the rotation and translation of each DgkA chain individually to their backbone beads for each 50 ns window. In the initial iteration, all backbone beads were used for alignment. In subsequent iterations, only backbone beads with RMSF < 0.1 nm were retained for fitting. This iterative fitting procedure was repeated until the list of rigid beads converged.^61,63^ The final RMSF profiles were obtained by averaging the backbone fluctuations across all 50 ns windows, chains, and replicas, resulting in one RMSF profile per photolipid conformation (STAPC/SCAPC).

#### 2.5.4 Photoswitchable membrane permeability

To study the impact of photolipid switching on membrane permeability, we simulated the membrane permeation of the `β`_2_ adrenergic receptor agonist salbutamol as well as the kinase inhibitor baricitinib. To estimate the free energy for drug permeation in a POPC bilayer with 25% D(T/C)APC, we calculated potentials of mean force (PMFs) using umbrella sampling (US). We first pulled the drug molecules from the water phase z = 4.3 nm above the membrane center to the other side of the membrane, i.e., z = −4.3 nm below the membrane center in order to generate the initial conformations for the US windows (force constant f_z_ = 1000 kJ/(mol nm^2^); pulling rate of −0.001 nm/ps). The reaction coordinate was defined as the z distance between the COM of the five beads composing the polar head of salbutamol,^63^ or the six initial beads of baricitinib corresponding to the bicyclic heteroaromatic core, and the COM of the terminal beads of the POPC tails (C4A and C4B). For the US, 85 windows were run for each photolipid conformation for each drug molecule from z = 4.2 nm to z = −4.2 nm. For both conformations, the windows had a distance of d_z_ = 0.1 nm and were sampled for 1 µs at 330 K. The reaction coordinate was restrained with a harmonic potential using a force constant of f_z_ = 1000 kJ/(mol nm^2^) for all windows. All PMFs were calculated using the GROMACS tool *gmx wham*.^64,65^ The initial 200 ns of each umbrella window were discarded as equilibration. Error estimation was done using a Bayesian bootstrap analysis with 100 bootstraps.

Drug-photolipid contacts. The analysis of the number of contacts of the drug molecules and D(T/C)APC lipids was performed employing the GROMACS tool *gmx select* together with a distance cutoff of 0.7 nm. For each window, the simulation was analyzed between 200-1000 ns in accordance with the PMF calculation.

**Figures.** Figures were prepared with the Python library Matplotlib,^66^ Gnuplot,^67^ and the program package VMD.

**Table 1:**
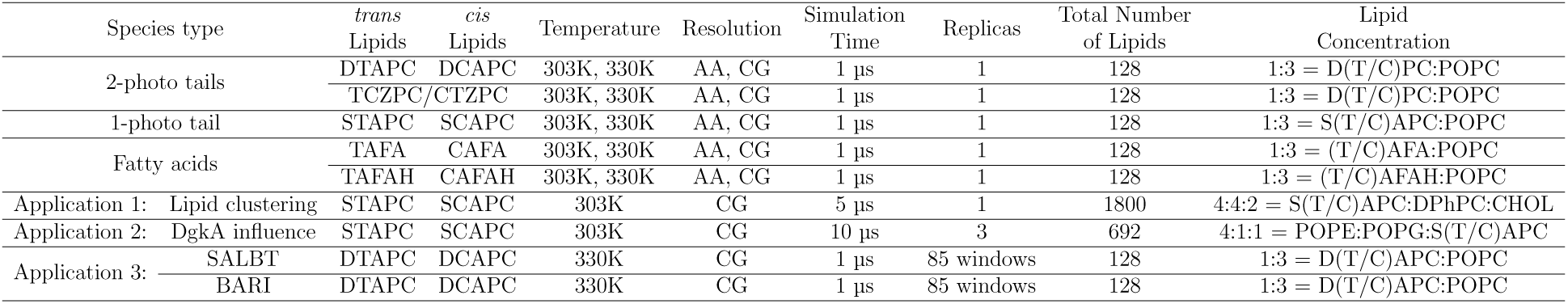
Overview of the photolipid simulations performed for this work.

## 3 Results and discussion

### 3.1 Azobenzene

The parametrization of the CG models for the azobenzene photoswitch was performed separately for the *cis* and *trans* isomers, following the strategies outlined by Alessandri et al..^51^ The mapping of azobenzene is shown in Figure 1 A. In line with the Martini 3 building block principle, each benzene ring in azobenzene was modeled using the established three-bead representation, where three TC5 beads are connected via constraints with LINCS^68^ with an equilibrium distance of 0.290 nm.

**Figure 1:**
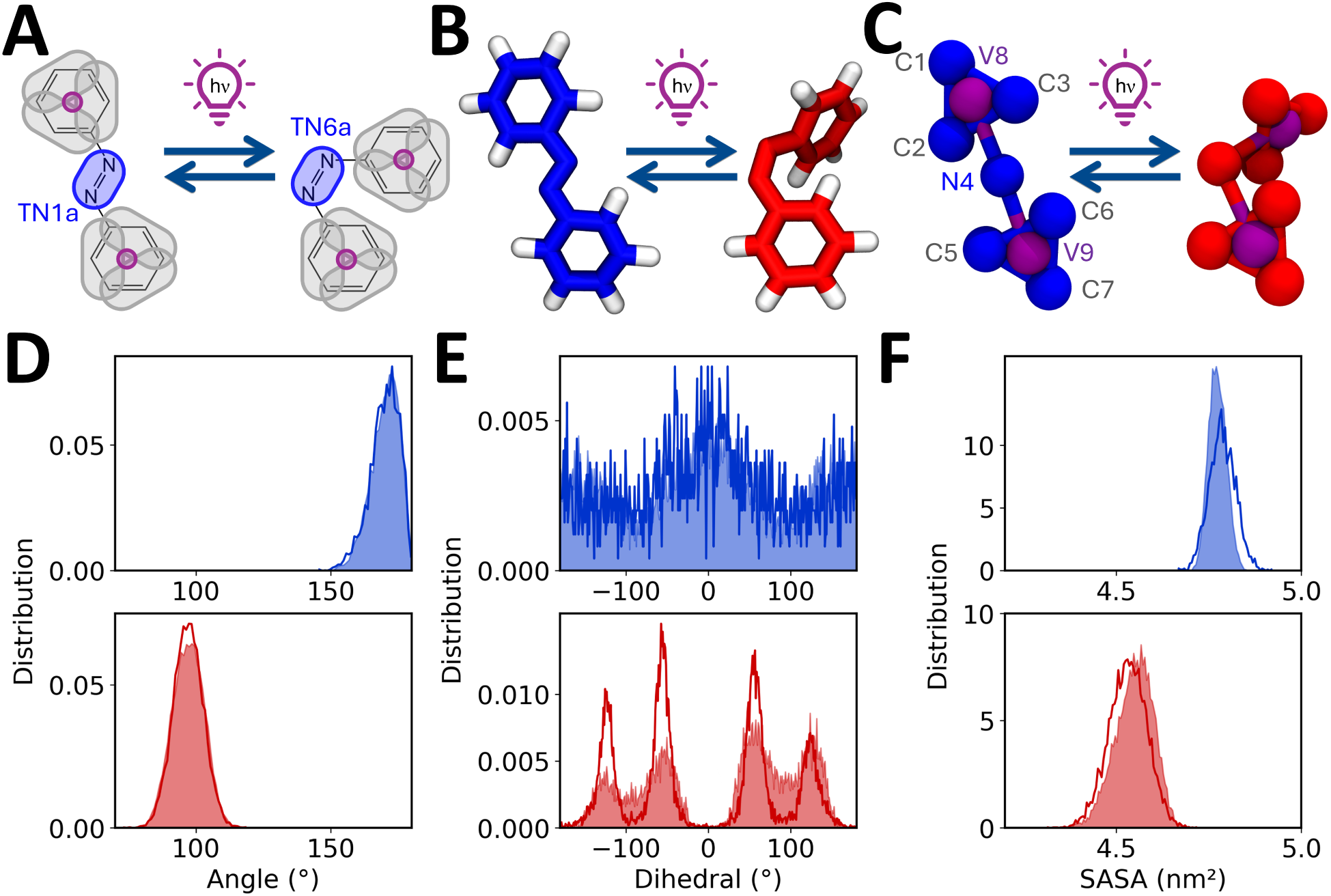
CG Martini 3 model for azobenzene. (A) CG Martini mapping of azobenzene. The bead type of N4 is indicated. All gray beads are assigned the bead type TC5; purple indicates virtual dummy beads. (B) Representative snapshots of the AA and (C) CG models visualizing the switching between *trans-* (blue) and *cis-*azobenzene (red). (D) Distributions of the crucial switching angle V8-N4-V9 obtained from pseudo-CG (mapped from AA, filled) and CG trajectories (line) for *trans-* (blue, upper panel) and *cis-*azobenzene (red, lower panel). (E) Distribution of the dihedral angle C2-C3-C7-C6 defining the relative orientation of the two phenyl rings; colors as in (D). (F) Solvent accessible surface area for *trans-* and *cis-*azobenzene.

Virtual dummy beads (V8 and V9) were placed at the center of each benzene ring and connected to the central azo bead (N4) via harmonic bonds with a force constant of k*_b_* = 60,000 kJ mol^−1^ nm^−2^. Four angles were introduced using ring beads (C2, C3, C5, C6), the respective dummy bead, and the central N4 bead. These prevent undesired rotation of the benzene rings around their central dummy beads. In addition, an angle potential between V8–N4–V9 was implemented to define the isomerisation state of the switch as shown in Figure 1D. A harmonic angle potential was used for the *cis-*isomer, while a linear angle potential was applied for the *trans-*isomer.

Four dihedral potentials were further introduced to both models. The dihedral angles C1–C2–C3–N4 and N4–C5–C6–C7 controlled the planarity of the azo moiety and the individual benzene rings. The C2–C3–C6–C5 dihedral determined the relative orientation between the two rings (Figure 1E). For the *cis-*isomer, additional dihedrals C2–V8–N4–V9 and C5–V9–N4–V8 were included to further regulate the orientation of each benzene ring relative to the central N4 bead. A comprehensive summary of all bonded interactions – distances, angles, and dihedrals – is provided in Figures S2–S4.

To evaluate the shape of the models, the SASA of both CG models was calculated. As shown in Figure 1F, the CG models accurately reproduce the difference in SASA between the two isomers and show excellent agreement with semi-empirical reference data.

Selecting an appropriate CG bead type for the central azo group (N4) posed a particular challenge due to the isomer-dependent dipole moment of the azobenzene photoswitch.^69^ To address this, the octanol–water partition coefficient (logP) was calculated and compared to experimental values. Based on this analysis, a TN1a bead was assigned to the *trans* isomer and a TN6a bead to the *cis* isomer. The resulting logP values (Table 2) reflect the experimentally observed trend: the *trans* isomer exhibits lower water solubility than the *cis* isomer due to its lower dipole moment. In a recently published CG Martini 3 model for azobenzene, the bead type TN3r was chosen for the azo-moiety and kept constant for both isomer.^70^ This results in a stronger deviation regarding the octanol-water patitioning coefficient (Table 2, right column). Moreover, the lack of dihedral angles in the alternative model leads to deviations in the ring positioning compared to atomistic reference data.

**Table 2:** Water-octanol partitioning of azobenzene.

|  | CG model | Brown et al.<sup>71</sup> | alt. CG model<sup>70</sup> |
| --- | --- | --- | --- |
| *trans*-azobenzene | 4.51 ± 0.03 | 4.20 | 3.71 ± 0.04 |
| *cis*-azobenzene | 3.33 ± 0.04 | 2.8 | 3.43 ± 0.03 |

The octanol-water partitioning behavior of the azobenzene isomers is consistent with their differential interactions with lipid bilayers, as shown in Figure 2. While both isomers of azobenzene integrate into the bilayer, the less polar *trans* isomer shows a higher density in the middle of the bilayer than the *cis* isomer, which interacts more with the polar lipid heads. This difference in membrane localization highlights the potential of azobenzene as a photoswitchable modulator of membrane properties.

**Figure 2:**
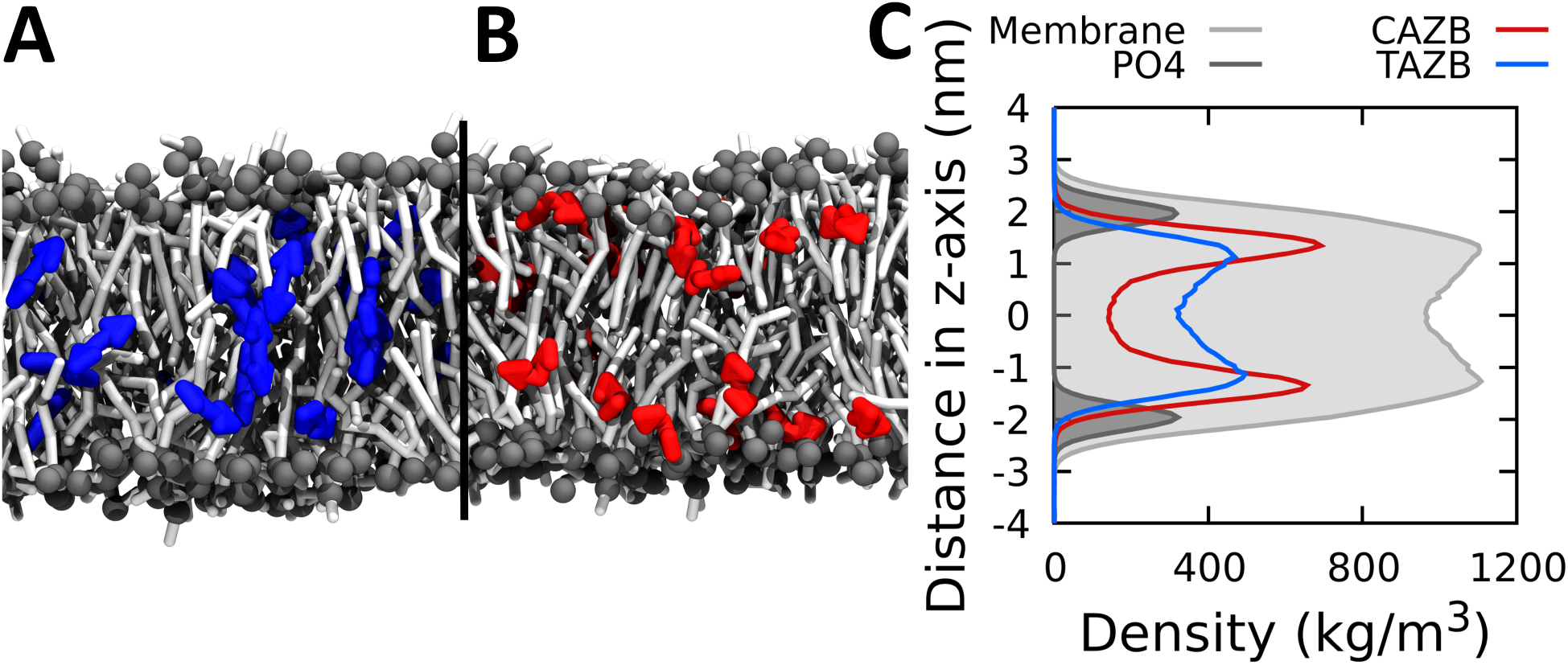
Interactions of azobenzene with a POPC membrane. (A,B) Final configurations of the *trans* (blue, A) and *cis* isomers (red, B) of azobenzene in a POPC membrane. (C) Mass densities of the whole membrane, phosphate group, and the azobenzene along the z axis. Azobenzene densities are scaled by a factor of 10.

### 3.2 Azobenzene-based photoswitchable lipids and fatty acids

Based on the azobenzene models, we developed CG Martini 3 models for two azobenzene-based photoswitchable lipids and a photoswitchable fatty acid at two protonation states. Figure 3A shows the currently most commonly used azobenzene-based PC lipid S(T/C)APC with one photoswitchable tail at the *sn2* position, which is commercially available. In our CG model, we connected the CG beads of azobenzene in *para* position of the azo moiety with the lipid tail beads. The three aliphatic C atoms towards the glycerol moiety are represented by a SC1 bead (bead name C1A), and the four aliphatic C atoms at the tail end are modeled as C1 bead (bead name C4A). The bonds between the lipid tail beads (C1A and C4A) and the azobenzene (C11 and C71) are adapted from the pseudo-CG trajectories generated from the AA simulations. As expected, the equilibrium bond lengths are shorter than in standard SC1-C1 and C1-C1 bonds due to the T beads of azobenzene. To capture the correct orientation and conformational rigidity of the azobenzene-containing tail, we introduced additional angles: GL1-C1A-C11, C1A-C11-V8, and V9-C71-C4A. Furthermore, the equilibrium angle and force constant of the angle GL2-GL1-C1A had to be modified to capture the correct tail conformations. Finally, the dihedral angle C1A-C21-C31-C11 was introduced to correctly capture the orientation of the first aromatic ring with respect to the lipid head group. The equivalent dihedral angle at the tail end, C71-C61-C51-C4A, was not necessary, and we omitted it for the sake of performance and stability of the CG model.

**Figure 3:**
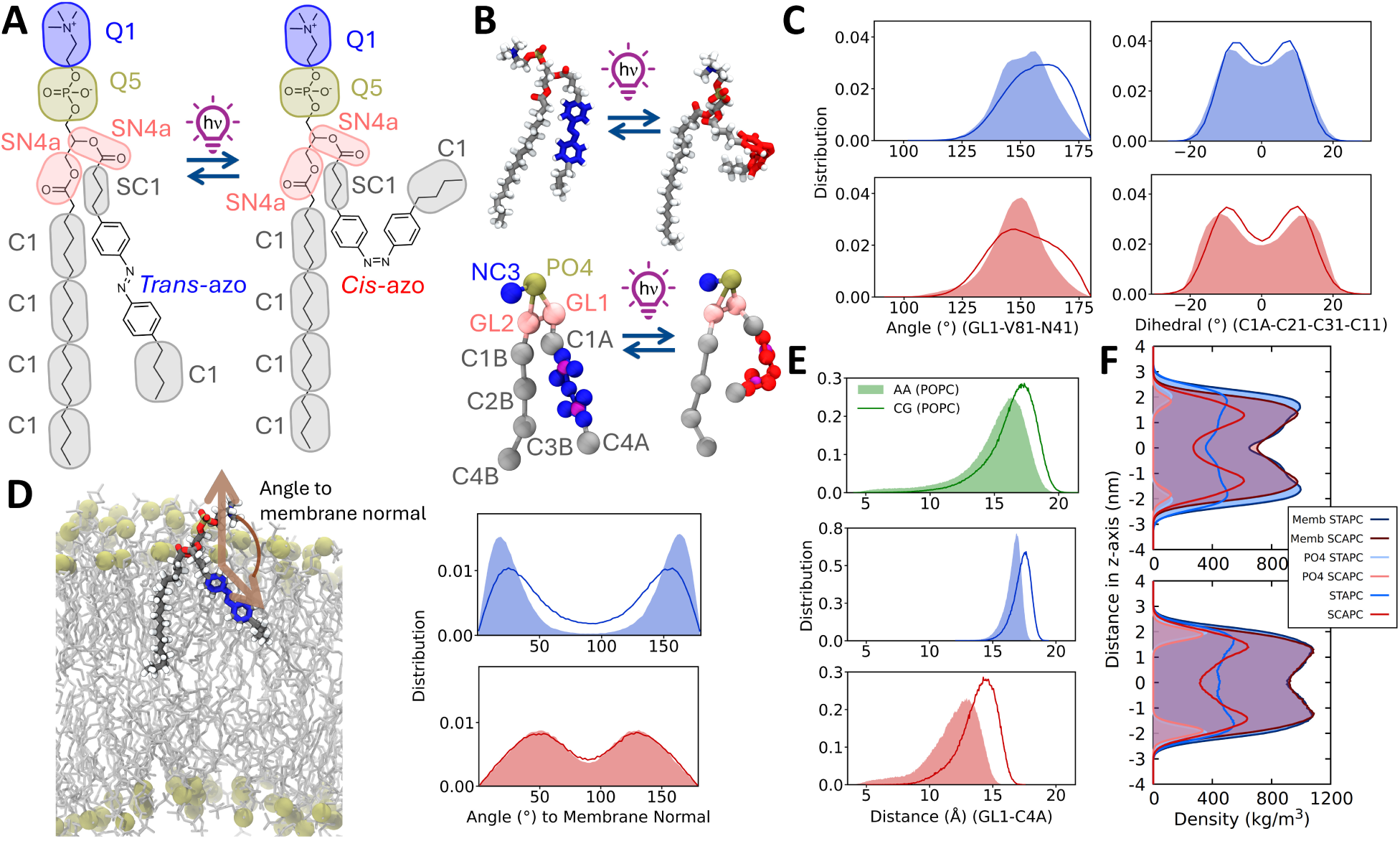
CG Martini 3 model for 1-stearoyl-2-(p-butyl-p-botanoyl-*trans-*/*cis-*azobenzene)-phosphatidylcholine (S(T/C)APC), the prototypical azobenzene-based photolipid. (A) CG Martini mapping of STAPC and SCAPC (including the bead types taken from the Martini 3.1 lipids). (B) Representative snapshots of the AA and CG models visualizing the switching between *trans-* (STAPC, blue) and *cis-*conformation of azobenzene (SCAPC, red). The CG *trans-*model includes the names assigned to the beads, which are the same in both models. (C) Distributions of exemplary bonded terms from pseudo-CG (mapped from AA, filled) and CG trajectories (line) visualizing the difference between both lipid states. The distributions show the angle GL1-V81-N41 along the photoswitchable tail starting from glycerol, and the dihedral angle C1A-C21-C31-C11 for the connection between azobenzene and the lipid tail, for the *trans* (blue, upper panels) and *cis* isomer (red, lower panel). (D) Angle distributions for the orientation of azobenzene with respect to the membrane normal for both lipid states (right). Schematic representation of the computed angle to the membrane normal (left). (E) Distributions of the GL1-C4A distance indicating the length of the photoswitchable lipid tail for the pseudo-CG (filled) and the CG trajectories. As a reference, the GL1-C4A distance distribution of POPC (green) is shown. (F) Mass densities of the whole membrane, phosphate group, and the photolipids S(T/C)APC for the AA (upper panel) and CG simulation (lower panel) along the z axis at 330 K. S(T/C)APC densities are scaled by a factor of 2.

Figure 3B shows representative snapshots of STAPC (left) and SCAPC (right) taken from bilayer simulations at atomistic (top) and CG resolution (bottom). They provide a visual impression of the overall shape and conformation of both *trans* and *cis* isomers. To quantify the similarity between the two resolutions and to validate the CG models of the photoswitchable lipids, we first compared key geometrical and conformational properties between AA (or pseudo-CG) and CG representations (Figure 3C–E and Figures S5–S14 in the SI). Figure 3C shows two exemplary distributions of the angle GL1-V8-N4 (left) and the dihedral angle C1A-C21-C31-C11 (right). Both show the close agreement between CG (lines) and pseudo-CG trajectories (filled) derived from the AA simulations. The distributions exhibit subtle differences between the *trans* and *cis* isomers of the photoswitchable lipid and confirm that the CG model is able to capture these differences. For example, the dihedral angle distribution (Figure 3C, right) exhibits a more pronounced central dip and broader distribution for the *cis* isomer compared to the *trans* isomer. The distributions for further bonded terms are shown in the Supporting Information (Figures S5-S14).

One crucial property of photoswitchable lipids is the orientation of the azobenzene moiety in the lipid tail with respect to the membrane normal, because it influences the changes in membrane order between both switching states. The CG model reproduces the angle between the vector spanned by the two aromatic ring centers and the membrane normal with good agreement for the *trans* isomer STAPC and with excellent agreement for the *cis* isomer SCAPC. This orientation is even better reproduced by the lipid model with two photoswitchable tails D(T/C)APC – particularly for the *trans* isomer (see Figure S11-S12). Thus, we hypothesize that the interaction with the second regular lipid tail might cause the deviations for the *trans* isomer STAPC in Figure 3D (right, top).

To compare the tail length between the two azobenzene conformations as well as the standard tails, we evaluate the distance between the glycerol backbone GL1 and the tail end C4A. The resulting distributions are depicted in Figure 3E. For STAPC, the maxima of the distributions are at 17.6 nm for the CG Martini 3 model and at 16.9 nm for the pseudo-CG data. These values change to 14.6 nm and 13.1 nm, respectively, after switching to the *cis* isomer SCAPC. Both differences are comparable with the difference for POPC between the CG Martini 3 (17.3 nm) and the pseudo-CG data (16.4 nm), which indicates a consistent overestimation of the tail length for the CG model (line) compared to the pseudo-CG data (filled). This is in line with a compromise made during the refined parametrization of the Martini 3 lipidome, where a small systematic decrease in the area per lipid was accepted to increase the reproduction of phase separation.^46^ This small area per lipid decrease entailed a slight overestimation of the membrane thickness. Figure 3E furthermore shows that the CG models reproduce well the widths of the pseudo-CG distance distributions. The slightly larger overestimation of the tail length in the case of SCAPC could be a starting point for future refinements of the CG models.

Figure 3F shows the mass density profiles along the membrane normal for the AA (top) and the CG model (bottom). The AA simulations show a reduction in membrane thickness upon isomerization, as expected, which becomes visible in the density of the entire membrane, as well as the densities of the phosphate groups. This trend is captured by CG simulations, although less pronounced. The smaller difference in membrane thickness may occur due to the more pronounced overestimation of the tail length for the *cis* isomer discussed above. The shape of the photolipid density matches well between AA and CG models, confirming that the CG model captures well the overall lipid arrangement in the membrane.

In addition to the photolipid with one photoswitchable tail, S(T/C)APC, discussed in detail above, we also parametrized a PC lipid with two photoswitchable tails, abbreviated D(T/C)APC, as well as the protonated and deprotonated photoswitchable fatty acids T/CAFAH and T/CAFA, which correspond to the photoswitchable tail before esterification.

The Martini building-block approach for the parametrization was followed across all species, ensuring consistent parameters among all *trans* and *cis* isomers of the photolipids. Thus, for the photolipid with two photoswitchable tails, the optimized parameters were transferred from the photoswitchable tail of S(T/C)APC to the second photoswitchable tail of D(T/C)APC. Slight modifications were required for the angles and dihedral angles involving the glycerol linker and the second tail to account for differences in the *sn1* and *sn2* position of the tail. In the case of the fatty acids, the angles along the tail were kept, but no dihedral angle was needed to keep the azobenzene orientation with respect to the lipid headgroup. Only an angle was required to correctly position the carboxyl/carboxylate group (COOH/COO bead). Distributions of the bonded terms as well as the densities and their comparison to pseudo-CG trajectories for all photolipids are depicted in Figures S5–S16 in the SI. They show the overall good agreement between our CG Martini 3 models and the pseudo-CG data obtained from AA reference simulations using CHARMM36.

To investigate the impact of the photolipids and their switching on a membrane, we simulated membranes composed of 25 mol% photolipid and 75 mol% POPC at 303 K and 330 K at both AA and CG resolution. The results presented in Figure 4 refer to the simulations at 330 K, which were used as the main reference to ensure that the membranes remain in the liquid phase during the simulation. The simulations at 303 K followed the same trends, and the results are provided in the SI (Figures S17). However, as the formation of a gel phase cannot be excluded,^23^ we will focus on the simulations at 330 K in the following. The membrane thickness was computed for each system as the average z-distance between P atoms (AA) and PO4 beads (CG), respectively, in the opposing leaflets (details see Section 2.4). For all photolipids at both resolutions, the *cis* isomer (Figure 4A–B, red) leads to a thinner membrane compared to the *trans* isomer (blue), which is due to a shortening of the photoswitchable tail and increased chain disorder. While this trend was well reproduced by the CG models (Figure 4B), the magnitude of the thickness difference is generally smaller than at AA resolution (Figure 4A). Figure 4C depicts the *trans-cis* difference at both resolutions. Nevertheless, the relative differences across the photolipid species are preserved by the CG models. In line with the decrease of membrane thickness upon *trans* to *cis* isomerization are the changes observed for the area per lipid (Figures 4D–F). The *cis* isomers occupy a larger area than the *trans* isomers, reflecting looser packing and increased lateral membrane expansion as the bilayer thins. Again, the CG models qualitatively reproduce this behavior (Figure 4E), slightly underestimating the differences relative to AA resolution (Figure 4D). Moreover, the switching also affects the properties of the non-photoswitchable lipid POPC at both resolutions (see Figure S18 in the SI).

**Figure 4:**
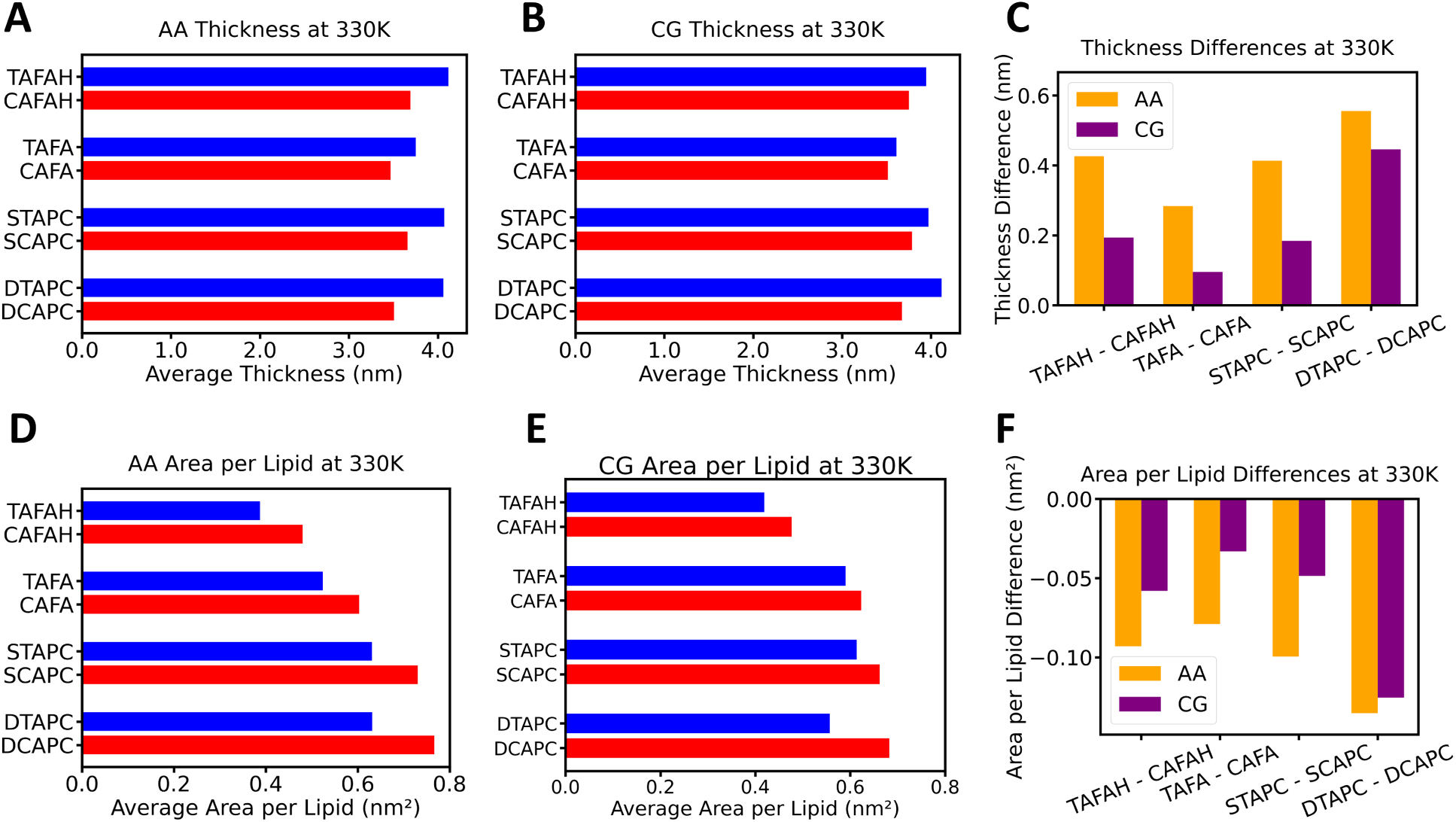
Properties for binary lipid mixtures containing 25 mol% azobenzene-based photolipid and 75 mol% POPC at 330 K. (A, B) Membrane thickness measured as the average z distance of the P atoms (AA, A) and of the PO4 beads (CG, B), respectively. The photolipids are protonated (CAFAH/TAFAH) and deprotonated fatty acid (CAFA/TAFA), PC with one photoswitchable tail (SCAPC/STAPC), and PC with two photoswitchable tails (DCAPC/DTAPC). (C) Thickness difference between *trans* and *cis* isomer of each photolipid species for AA (orange) and CG systems (purple). (D, E) Area per lipid of the photolipids for the AA (D) and CG systems (E). (F) Area per lipid difference between *trans* and *cis* isomers of each photolipid species for AA (orange) and CG systems (purple).

Among all systems, the largest difference upon isomerization was observed for the lipid with two photoswitchable tails, DTAPC and DCAPC, respectively. This can be explained by an additive effect of incorporating two photoswitchable tails instead of one, which causes a larger modulation of the lipid organization and membrane properties upon isomerization. In contrast, the deprotonated photoswitchable fatty acid (T/C)AFA produces the smallest effect upon isomerization.

These trends are identical for both resolutions, highlighting that our CG Martini 3 models successfully reproduce the relative behavior at AA resolution.

In contrast to S(T/C)APC, D(T/C)APC, and (T/C)AFA, which all have a head group containing charged beads, the protonated fatty acid (T/C)AFAH instead has a polar P2 bead as head group. While the polar bead still favorably interacts with the water phase, the energetic penalty for entering the hydrophobic core of the membrane is much lower than for charged Q beads. This impacts the behavior of (T/C)AFAH in comparison to the other photolipids parametrized here. In case of (T/C)AFAH, the change in polarity from *trans-*to *cis-*azobenzene is decisive enough to allow a higher population of TAFAH than CAFAH in the inter-leaflet space. This increases the membrane thickness for the *trans* compared to the *cis* isomer. This contribution to the modulation of membrane properties originates from a different molecular mechanism than in the case of the photolipids with charged headgroup beads. Representative snapshots are depicted in Figure S19 in the SI.

### 3.3 Application examples

#### 3.3.1 Controlling membrane domain formation

Following the parametrization and successful comparison to AA reference simulations, we applied our new CG photolipid parameters in three test cases of higher complexity to showcase their diverse potential applications. The first test case is a ternary lipid mixture for which photocontrol of domain formation has been demonstrated experimentally in giant unilamellar vesicles.^14^ To this end, we simulated the ternary lipid mixture DPhPC:STAPC:cholesterol at a 4:4:2 ratio for 5 µs. Figure 5 (top) shows snapshots of the membrane patch at different simulation times. At the beginning, the lipids are well mixed (top left), and within approximately 500 ns, two domains are formed, consisting of STAPC/cholesterol and DPhPC, respectively. The second snapshot shows the stable domains after 5 µs of simulation. At this point, we modeled the instantaneous photoisomerization by switching the CG model from STAPC to SCAPC. After a brief equilibration (50 ps with timestep of Δt = 1 fs), we continued the simulation for additional 5 µs. The third snapshot in Figure 5 (top) shows that the domains dissolve quickly within approximately 500 ns. After 5 µs, the system exhibits a stable fully mixed ensemble.

**Figure 5:**
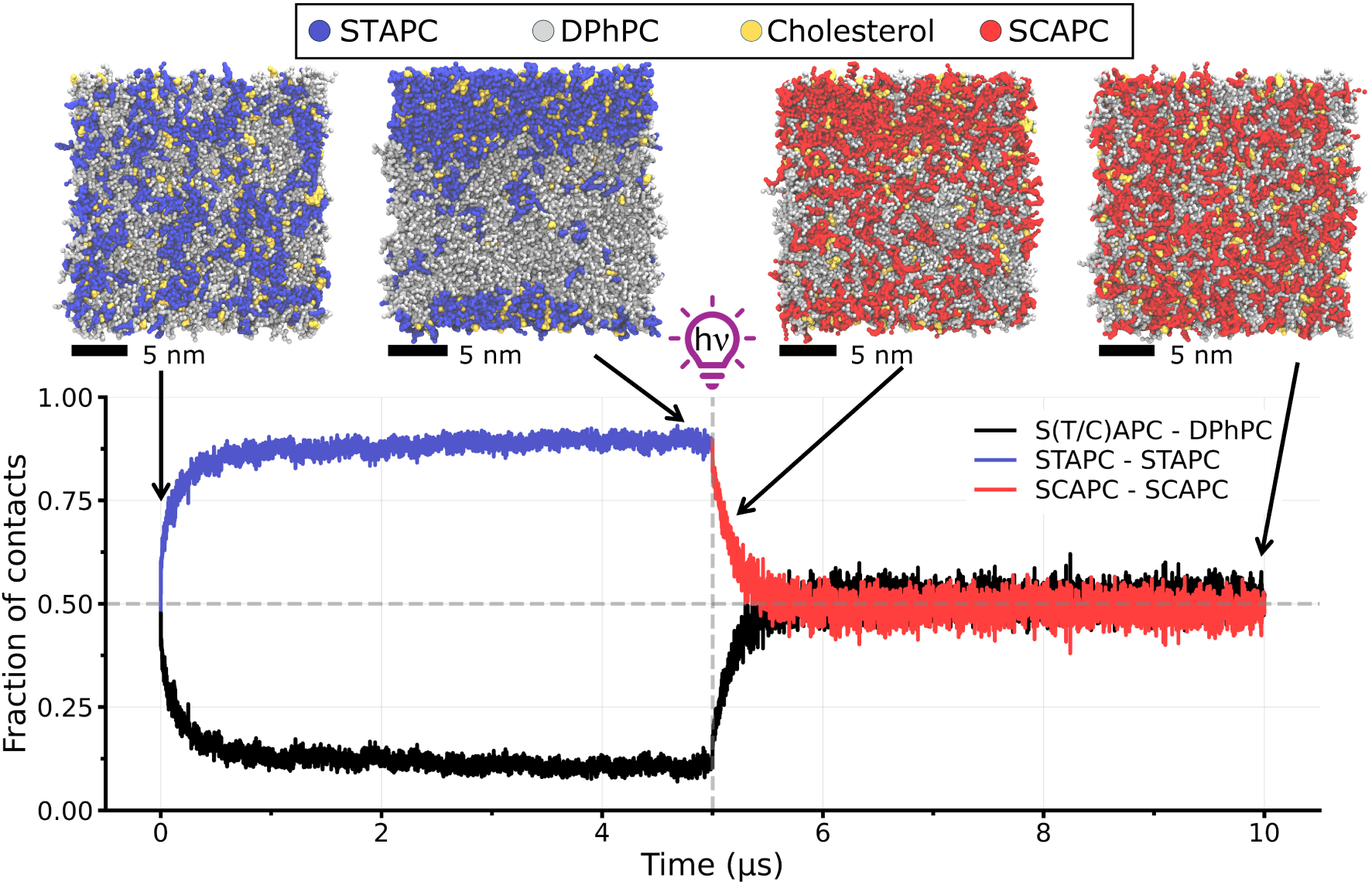
Light-controlled domain formation in the ternary lipid mixture S(T/C)APC:DPhPC:cholesterol = 2:2:1. The top row shows snapshots of the membrane patch taken after 1 ns, 5 µs, 6 ns after the switching, and 5 µs after the switching. The two snapshots on the left show the system with STAPC (*trans-*azobenzene conformation) in the random starting configuration (1 ns) and after domain formation (5 µs). The two snapshots on the right show the system 6 ns after switching the CG photolipid model to SCAPC (*cis-*azobenzene conformation), followed by the dissolution of the two domains (5 µs after switching). The bottom graph shows the time evolution of the fraction of contacts between the phospholipids S(T/C)APC and DPhPC for the total simulation time of 10 µs.

Figure 5 (bottom) shows the contact fraction between the photolipids and DPhPC, quantifying the domain formation. Starting from a homogeneous mixture, with STAPC-STAPC and STAPC-DPhPC contact fractions being around 0.5, the values rapidly evolve to ap-proximately 0.9 and 0.1, respectively. This trend is in line with the clear domain formation observed in the snapshots. Upon switching to the *cis* isomer SCAPC, the contact fractions – now SCAPC-SCAPC and SCAPC-DPhPC – quickly revert to approximately 0.5, indifcating the reformation of a homogeneous mixture.

These results nicely demonstrate that our CG Martini 3 model is able to capture the different domain-forming properties of both isomers, which is consistent with experimental observations.^14^

As a second case for light-controlled domain formation, we simulated the binary lipid mixture D(T/C)APC:POPC at a 1:3 ratio for 5 µs at 330 K. Vesicles with D(T/C)APC have been used experimentally to study their applicability for photocontrolled drug delivery.^15,16^ Figure 6 (top) shows snapshots of the membrane patch at different simulation times. Initially, the bilayer is well mixed (top left). Within the first few hundred nanoseconds, a DTAPC domain is formed. The second snapshot shows the stable domain after 4.8 µs of simulation. At this point, the instantaneous photoisomerization was modeled by switching the CG model from DTAPC to DCAPC. After a brief equilibration similar to the ternary mixture, the simulation was continued for another 5 µs. The third snapshot shows that the domains dissolve quickly within a few hundred nanoseconds, and after 4.8 µs, the system is fully mixed (rightmost snapshot).

**Figure 6:**
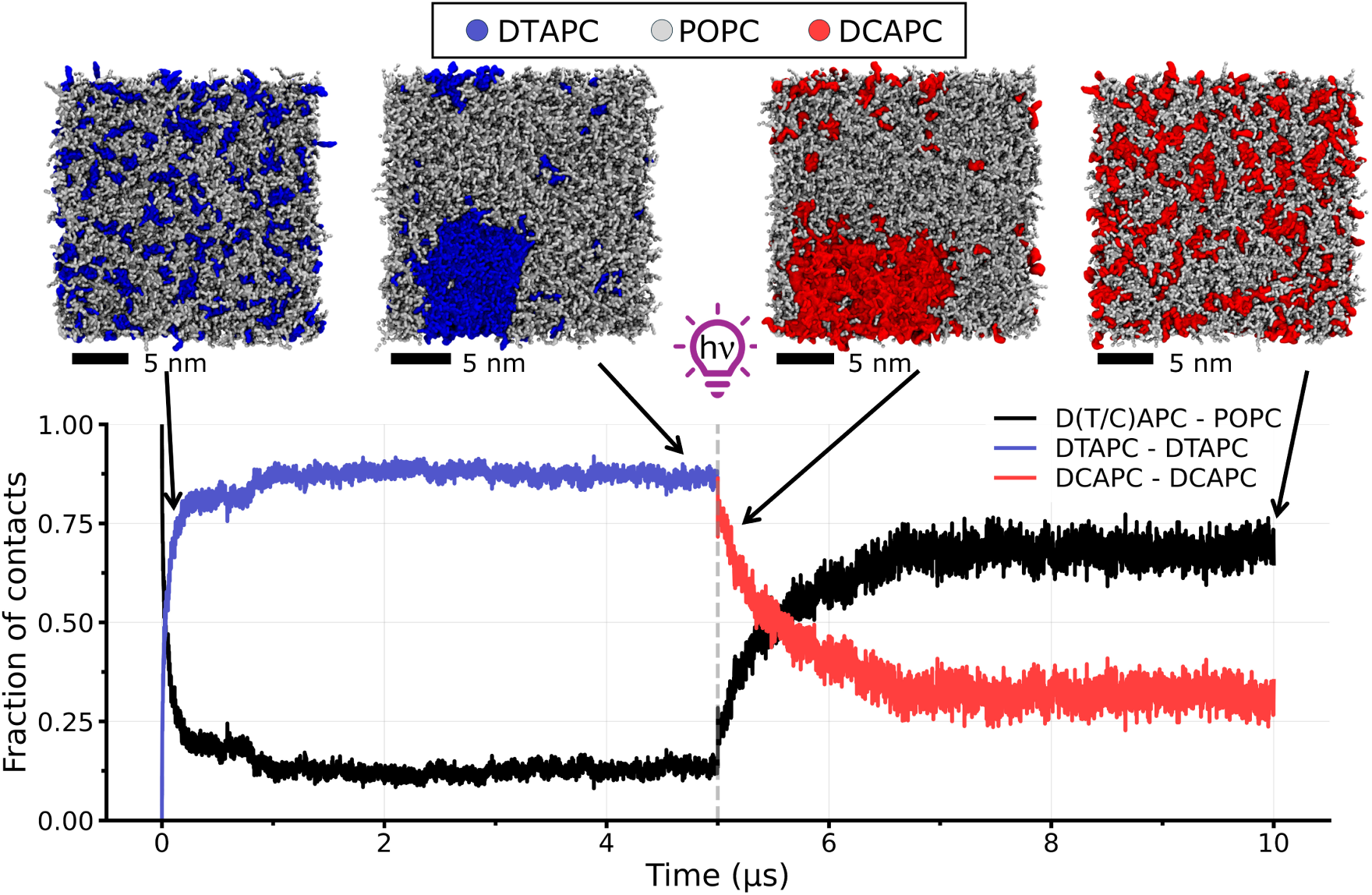
Light-controlled domain formation in the binary lipid mixture D(T/C)APC:POPC = 1:3. The top row shows snapshots of the membrane patch taken after 100 ns, 4.8 µs, 100 ns after the switching, and 4.8 µs after the switching. The two snapshots on the left show the system with DTAPC (*trans-*azobenzene conformation) in the random starting configuration (100 ns) and after domain formation (4.8 µs). The two snapshots on the right show the system 100 ns after switching the CG photolipid model to DCAPC (*cis-*azobenzene conformation), followed by the dissolution of the two domains (4.8 µs after switching). The bottom graph shows the time evolution of the fraction of contacts between the phospholipids D(T/C)APC and POPC for the total simulation time of 10 µs.

The time evolution of contact fractions between D(T/C)APC and POPC is shown in Figure 6 (bottom). Starting from a homogeneous mixture, DTAPC–DTAPC and D(T/C)APC–POPC contact fractions rapidly evolved to approximately 0.9 and 0.1, respectively, indicating robust domain formation as for the ternary mixture. After switching to the *cis* isomer DCAPC, these contact fractions rapidly returned to values near 0.3 and 0.6, consistent with complete mixing for this membrane composition.

#### 3.3.2 Regulating the flexibility of DgkA

In the second test case, we studied the influence of a photoswitchable lipid membrane on the dynamics of embedded transmembrane proteins. Doroudgar et al. demonstrated experimentally by means of solid-state NMR measurements that a small but measurable effect is observable for the DgkA trimer.^18^ To investigate whether our CG photoswitchable lipid models induce the changes in CG protein flexibility upon *trans-cis* isomerization, we simulated the trimeric transmembrane protein DgkA embedded in a photoswitchable membrane with the experimental lipid composition of POPE:POPG:S(T/C)APC at a ratio of 24:6:5 (Figure 7A). We performed three replicas of 10 µs for each photolipid isomer, resulting in a total sampling of 90 µs (3×3×10 µs) for a DgkA monomer. Figure 7B depicts the backbone RMSF averaged over the three chains and three replicas. Overall, the RMSF profiles are very similar for both photolipid isomers in line with the experimental data.^18^ A closer look reveals an increase in flexibility for the first 25 N-terminal residues of DgkA in the SCAPC-containing membrane (red line, see also inset). These residues form the helix positioned at the membrane surface and interact closely with the lipid headgroups. This might render them sensitive to the *trans-cis* isomerization, modulating the membrane properties. The three transmembrane helices show virtually identical RMSF for both lipid environments, indicating that the core of the protein does not sense the photoswitching. To visualize the difference in surface helix flexibility, Figure 7C shows the conformational ensemble of the surface helices obtained from one replica (for all replicas, see Figure S20). While both systems preserve the characteristic trimeric structure of DgkA, the SCAPC system (right) displays an increased mobility of the surface helix, consistent with the increased lipid disorder induced by the *cis*-photolipid. These results suggest that light-controlled membrane composition may cause local, tunable effects on protein flexibility. The increased flexibility observed in our simulation upon isomerisation is in the same region as the most flexible residues identified experimentally.^18^ This agreement highlights the ability of the Martini 3 model to reproduce the experimentally reported modulation of membrane protein flexibility by photolipids. There might still be room for improvement in capturing the protein dynamics more precisely, because the protein model of DgkA was not further optimized. This optimization might enable an even clearer differentiation between the *trans-* and *cis-*photolipid environments. In particular, the a GōMartini 3 protein model was used with the default parameters to maintain the native fold of DgkA. Optimizing the Gō-like potentials may allow a more sensitive assessment of the protein’s response to the membrane environment.

**Figure 7:**
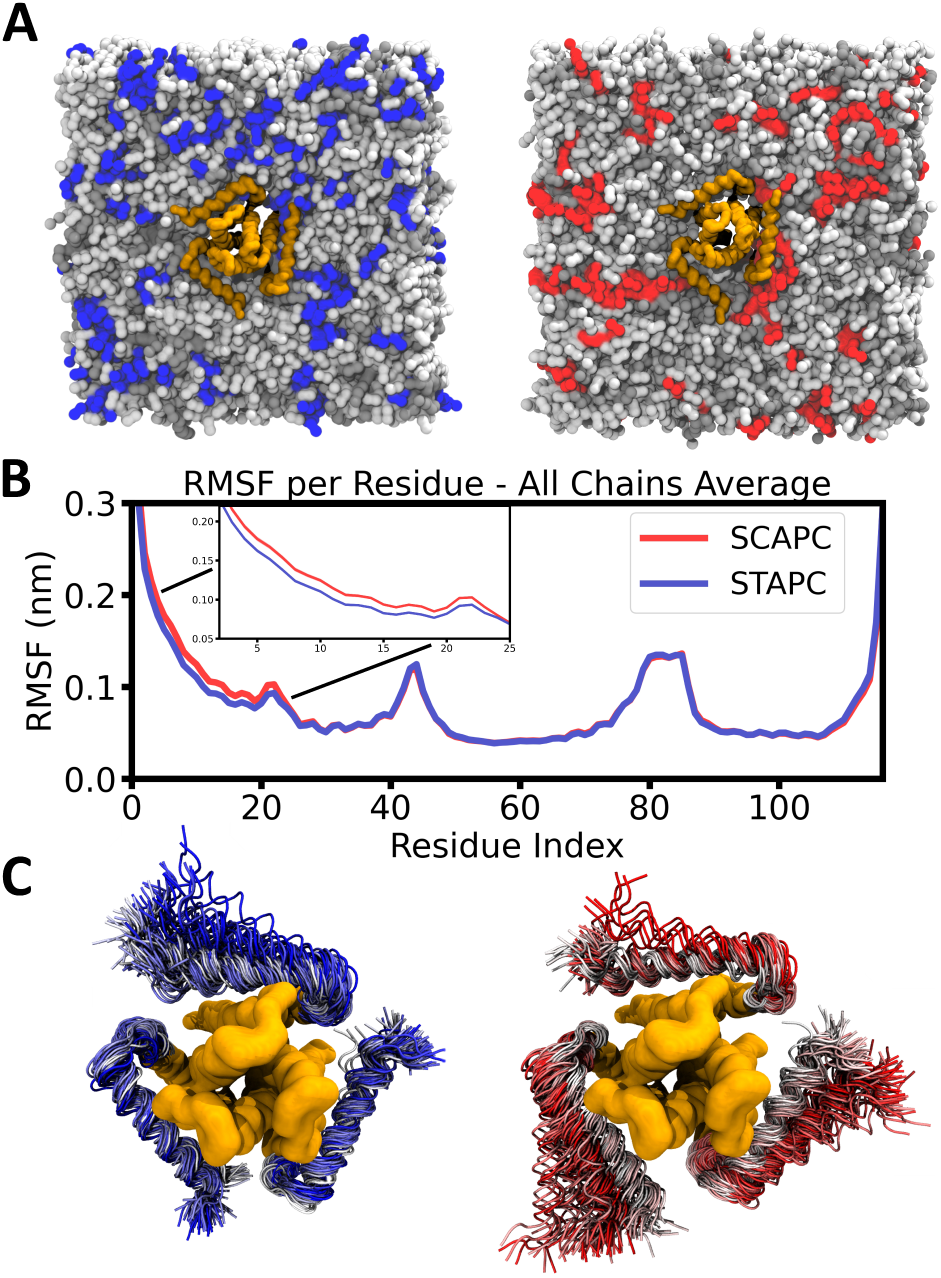
Light-controlled modulation of the flexibility of the transmembrane protein DgkA embedded in a photoswitchable lipid membrane. (A) Example snapshots of DgkA trimers (light orange) in the membrane patch composed of POPE (white), POPG (grey), and STAPC (blue) or SCAPC (red), respectively, at a ratio of 24:6:5. (B) Backbone root mean square fluctuation (RMSF) of DgkA embedded in STAPC- and SCAPC-doped membranes. Photoswitching to the *cis* isomer of the lipids leads to a flexibility increase of the surface helix of DgkA (residues 1-25). (C) Movement of the surface helix of a DgkA trimer in a STAPC-(left) and SCAPC-containing (right) membrane. 100 equally spaced frames of a 10 µs trajectory are depicted. All replicas are shown in Figure S20 in the SI.

#### 3.3.3 Enhancing membrane permeation

Liposomes containing photolipids are promising candidates for photocontrolled drug delivery, because the *trans-cis* isomerization of the photolipid enhances membrane permeability and thereby enables light-controlled drug release.^7,15–17^ In the third test case, we wanted to assess whether our CG Martini 3 photolipid models capture changes in membrane permeability and calculated the free energy barrier for membrane permeation of salbutamol and baricitinib, an agonist of the G protein-coupled receptor `β`_2_ adrenergic receptor and a kinase inhibitor, respectively. To this end, we pulled a salbutamol or baricitinib molecule across two membrane patches composed of POPC and DTAPC (Figure 8A and Figure 9A)/DCAPC (Figure 8B and Figure 9B) at a ratio of 3:1. From these pulling simulations, we extracted configurations along the membrane normal from 4.2 nm above to 4.2 nm below the membrane center, and conducted US simulations to calculate the PMF and to quantify the energy difference for membrane permeation. Each window was sampled for 1 µs. Figure 8C and 9C depict the resulting PMFs for the *trans* isomer DTAPC (blue) and the *cis* isomer DCAPC (red).

**Figure 8:**
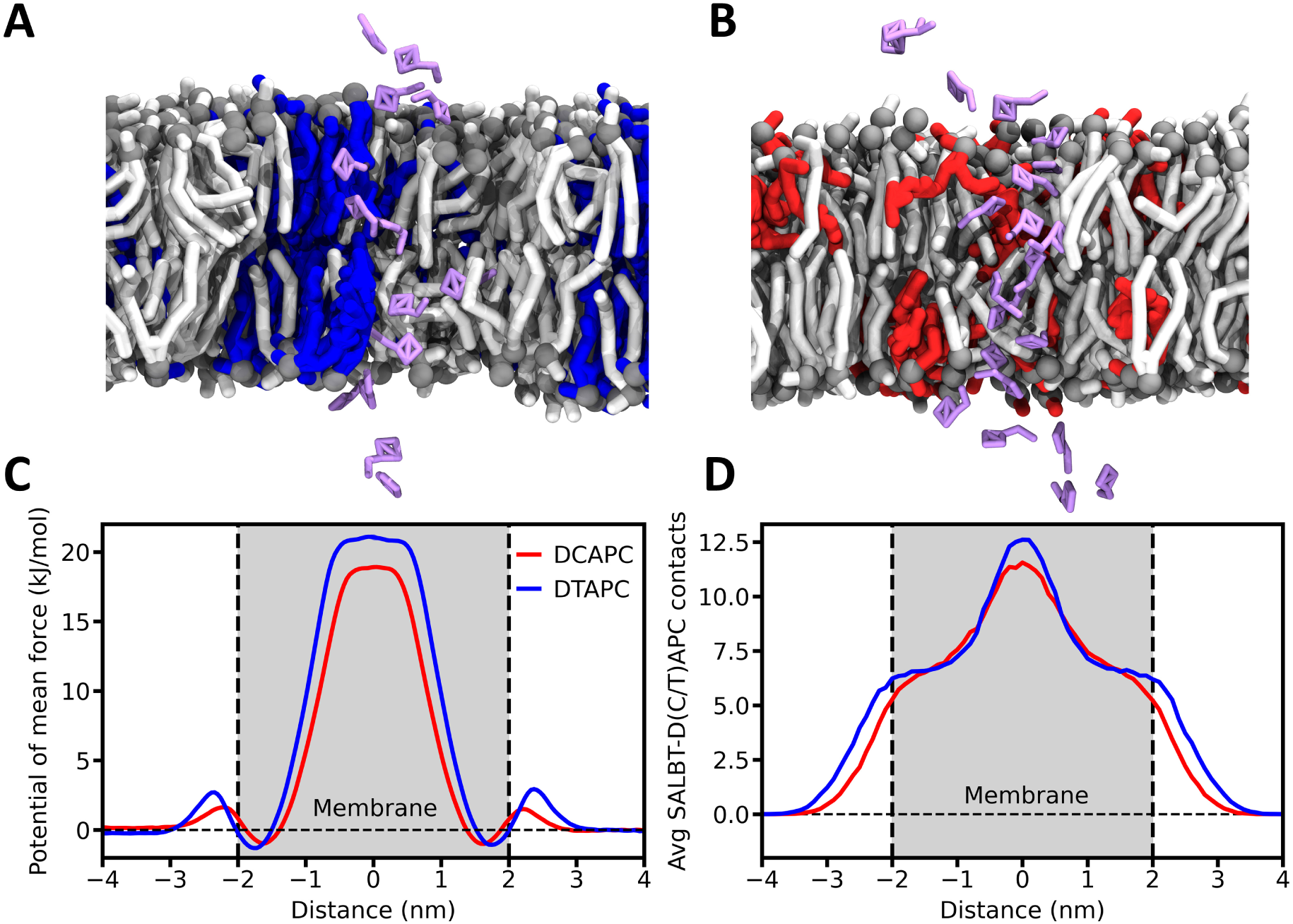
Free energy calculations for the permeation of salbutamol through the photo-switchable lipid membrane D(T/C)APC:POPC = 1:3. (A,B) Initial configurations of the US windows extracted from a pulling simulation of salbutamol (purple) along the membrane normal with DTAPC (blue, A) and DCAPC (red, B). (C) PMFs for the membrane permeation of salbutamol along the membrane normal calculated using US. Error bars are below the line width. (D) Average number of contacts between salbutamol and D(T/C)APC per US window along the reaction coordinate. Error bars are below the line width.

**Figure 9:**
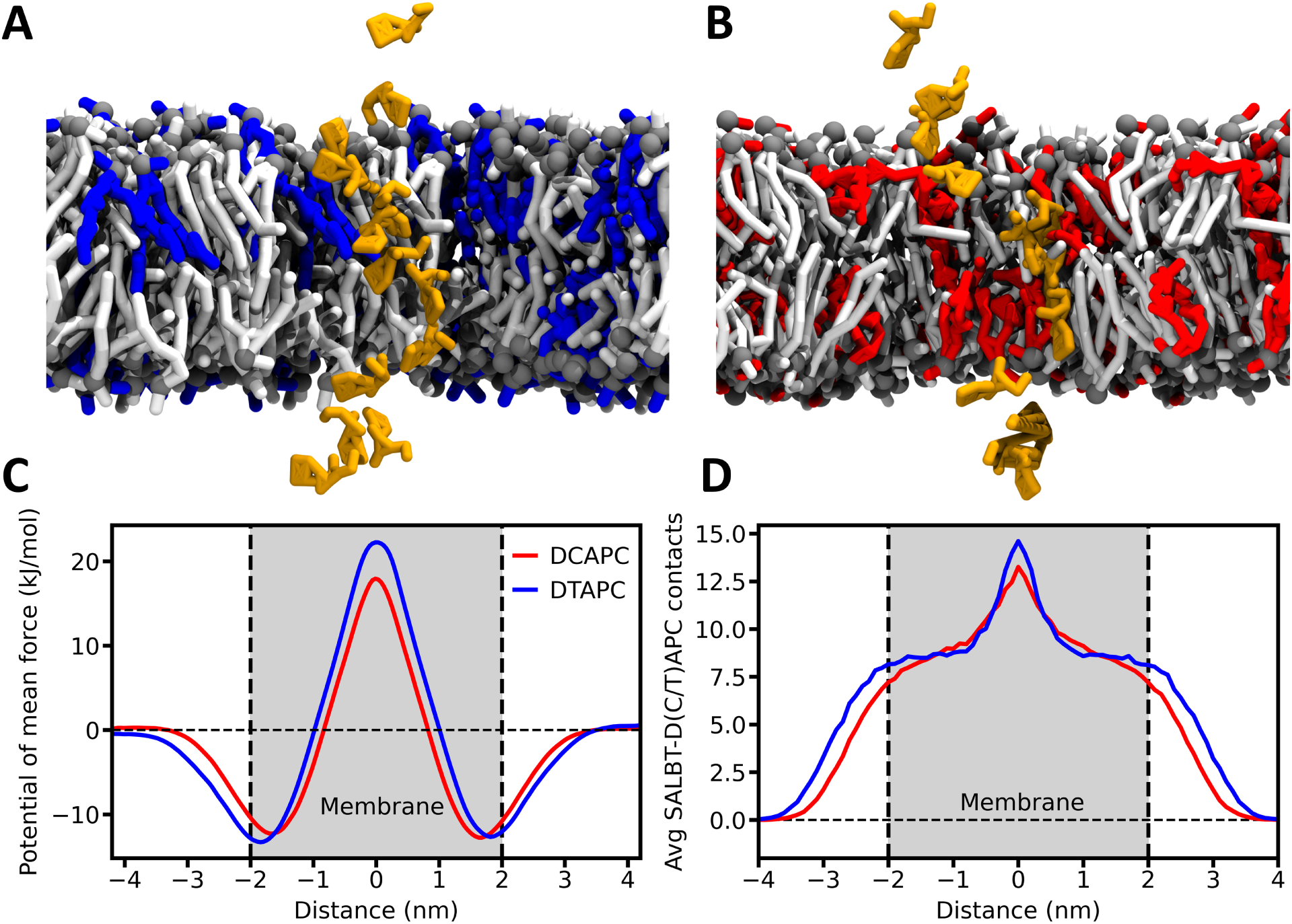
Free energy calculations for the permeation of baricitinib through the photoswitchable lipid membrane D(T/C)APC:POPC = 1:3. (A,B) Initial configurations of the US windows extracted from a pulling simulation of baricitinib (yellow) along the membrane normal with DTAPC (blue, A) and DCAPC (red, B). (C) PMFs for the membrane permeation of baricitinib along the membrane normal calculated using US. Error bars are below the line width. (D) Average number of contacts between baricitinibg and D(T/C)APC per US window along the reaction coordinate. Error bars are below the line width.

The free energy barrier for salbutamol permeation through the DTAPC-containing membrane from the water phase is approximately 21.1±0.2 kJ/mol, while it decreases to around 18.8±0.1 kJ/mol for the DCAPC-containing membrane. We estimated the uncertainty of the PMFs as the free energy difference in water on both sides of the membrane, averaging over the regions from 3.2 to 4.0 nm and from -3.2 to -4.0 nm, which is 0.2 kJ/mol or smaller for both systems. This estimate, however, is larger than the error estimated by the WHAM analysis, but describes the sampling error better in our view. The free-energy barrier was calculated as the difference between the average free energy in the membrane center (–0.2 to 0.2 nm) and the average free energy in bulk water (3.2 to 4.0 nm and –3.2 to –4.0 nm). The effect of the *trans-cis* isomerization on the barrier height aligns qualitatively with experimental findings on liposome permeability.^7,15–17^ The lower energy barrier in the *cis-*membrane is likely due to increased membrane disorder upon isomerization, which can facilitate membrane permeation.^72–74^ To examine whether salbutamol samples regions of similar lipid compositions in both switch states, we calculated its average number of contacts with D(T/C)APC along the reaction coordinate. The results, depicted in Figure 8D, show that salbutamol interacts similarly with both photolipids throughout the membrane except for the membrane center, where it forms slightly more contacts with DTAPC. We attribute this higher number of contacts to the more extended conformations of DTAPC compared to DCAPC (see Figure 3E) and not to a sampling of different lipid compositions. This conformational difference is also reflected in the density distributions along the membrane normal (Figure S15-S16 in the SI), and gives salbutamol the possibility to be in contact with more DTAPC molecules than DCAPC ones in the membrane center. Moreover, the increased membrane thickness for DTAPC is also visible in the salbutamol-D(T/C)APC contacts.

The free energy barrier for baricitinib permeation through the DTAPC-containing membrane from the water phase is approximately 22.2±0.9 kJ/mol, while it decreases to around 17.8±0.1 kJ/mol for the DCAPC-containing membrane. We estimated the uncertainty of the PMFs as the free energy difference in water on both sides of the membrane, averaging over the regions from 3.5 to 4.0 nm and from -3.5 to -4.0 nm, which is 0.9 kJ/mol or smaller for both systems. As before, this estimate is larger than the error obtained by the WHAM analysis. The free-energy barrier was calculated as the difference between the average free energy in the membrane center (–0.1 to 0.1 nm) and the average free energy in bulk water (3.5 to 4.0 nm and –3.5 to –4.0 nm).

The results, depicted in Figure 9D, show that baricitinib interacts similarly with both photolipids throughout the membrane except for the membrane center, where it forms slightly more contacts with DTAPC, similar to salbutamol.

The difference in barrier height of 2.3±0.2 kJ/mol for salbutamol and 4.4±0.9 kJ/mol for baricitinib corresponds to a reduction of around 10–20%, which qualitatively agrees with reported changes from AA simulations. For doxorubicin, changes in barrier height of around 10 kJ/mol were reported^23^ for an overall barrier height of around 65 kJ/mol with the *trans* isomer in a system containing 14 mol% STAPC. For phenol and a dopamine D1-like receptor agonist, changes in barrier height of about 5.4 kJ/mol and 7.0 kJ/mol, respectively.^25^ In the latter study, however, the membrane was composed exclusively of azoPC, and for the *trans* isomer, it was in the gel phase. Moreover, limited sampling at AA resolution might be the reason for the large difference in overall barrier height for doxorubicin compared to the molecules studied here.^75^ The two examples show that our CG Martini 3 models for azobenzene-based photolipids provide a framework for studying light-controlled membrane permeation and could, with further validation, be extended toward more systematic screenings of lipid compositions for photocontrolled drug release.

### 3.4 Limitations

While the CG Martini 3 models of azobenzene and azobenzene-based photolipids capture the conformational differences between *trans* and *cis* isomers remarkably well, they entail some limitations, which we will discuss in this section. The CG models reproduce the (meta)stable conformations of the azobenzene moiety in the electronic ground state, but they are not suitable to study its ultrafast switching process. This switching process starts in the electronically excited state, which requires quantum mechanical treatment of the molecules. The distinct models can, however, be used to study the impact of photoswitching on the reorganization of a photoswitchable membrane on the ns–µs time scale as shown in Section 3.3.1. Furthermore, the difference in dipole moment of the two isomers is effectively modeled by using a different bead type (*trans* – TN1a; *cis* – TN6a; see Figure 1). This assignment bends the rules for parametrization in Martini 3, but azobenzene is a special case in this regard due to the change of the molecular symmetry during switching. We expect that substitutions on the aromatic rings, which significantly impact the electron density on the azo moiety, might not follow the fragment building block approach of Martini 3 and require further bending of the parametrization rules.

An additional limitation relates to the all-atom reference model used for the azobenzene moiety. The parameters from Klaja et al. provide the most thoroughly characterized quantum chemically–derived description of azobenzene in a lipid-like context, specifically in a fatty acid. In our workflow, only the azobenzene fragment was taken from this study, while the remaining lipid components were directly adopted from the CHARMM36 lipid force field. As a result, the CG models accurately capture the conformational behavior of the azobenzene moiety and reproduce experimentally observed membrane phenomena. Nevertheless, future experimental reference data for azobenzene-containing lipids could enable refinement of the all-atom parameters, which in turn would allow systematic re-optimization of the CG models.

A limitation with regard to the time scale in CG MD simulations which also applies here is that the effective CG time scale does not correspond to real time due to the smoothening of the potential energy surface.^61^ Concerning the impact of photolipid switching on embedded transmembrane proteins, we recommend a careful parametrization of the CG protein model in order to capture the impact as good as possible.^58,61^ Forthcoming improvements of the Martini 3 protein model will potentially improve this aspect further.

## 4 Conclusions

We presented CG Martini 3 models for azobenzene and azobenzene-based photoswitchable lipids. Overall, the trends within different photolipids parametrized in this work are in semi-quantitative agreement with the AA reference simulations. Moreover, we showed that experimental findings can be reproduced, such as photocontrol of lateral phase separation in lipid mixtures, the modulation of protein flexibility upon photoswitching, as well as changes in membrane permeability. We therefore expect our Martini 3 models of azobenzene-based photolipids to be useful for the community to study the impact of photolipids in large biomolecular ensembles at almost atomistic resolution.

In the future, we aim to extend the set of available azobenzene substitution patterns, which will open up further applicability of the CG models in the field of photopharmacology and material science.

## Acknowledgement

The authors thank the Alfons und Gertrud Kassel Foundation, the Dr. Rolf M. Schwiete Foundation, the H. & E. Kleber Foundation, the Center for Multiscale Modelling in Life Sciences (CMMS) sponsored by the Hessian Ministry of Science and Art, and the SCALE cluster of excellence initiative for funding. Furthermore, the authors thank the Center for Scientific Computing at Goethe University Frankfurt for access to FUCHS and Goethe-HLR.

## Supporting Information Available

The Supporting Information is available free of charge at https://pubs.acs.org/doi/….

Details of the parametrization of the CG models, membrane properties, DgkA flexibility, and the PMF convergence (PDF).

## TOC Graphic

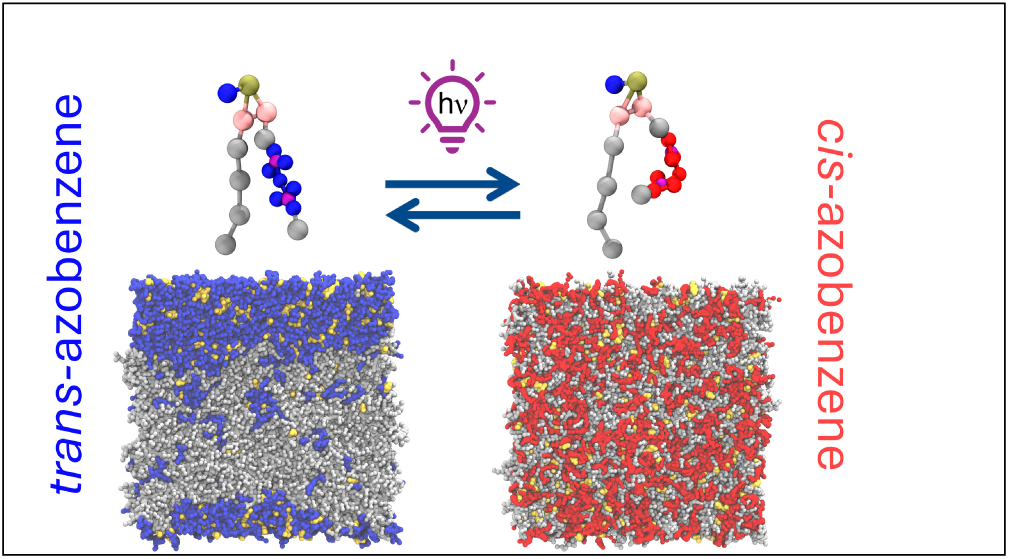

