## Supporting Information for "Martini 3 coarse-grained models of azobenzene-based photolipids: Modulation of membranes with light"

### Supporting Information File: Martini 3 coarse-grained models of azobenzene-based photolipids: Modulation of membranes with light

Cristina Gil Herrero,[†,‡] Thilo Duve,[†,‡] and Sebastian Thallmair[*,†]

†*Frankfurt Institute for Advanced Studies, Ruth-Moufang-Str. 1, 60438 Frankfurt a. M., Germany*

‡*Goethe University, 60438 Frankfurt a. M., Germany*

#### Contents

**1 Thermodynamic Integration** **3**

**2 Azobenzene Parametrization** **3**

**3 Bonded terms** **6**

**4 Density plots** **16**

**5 Membrane thickness and area per lipid analysis** **18**

**6 Comparative snapshots of (T/C)AFAH and S(T/C)APC** **20**

**7 DgkA flexibility of the three replicas** **21**

**8 US convergence tests and histograms** **21**

**References** **23**

### 1 Thermodynamic Integration

We used thermodynamic integration to compute the solvation free energies of azobenzene in water (W) and hydrated octanol. To simulate hydrated octanol, approximately 0.3 mole fraction of water was added to the simulation box.[1] For each solvent–solute combination, 21 simulations were performed with $\lambda$ points equally spaced between 0 and 1. Lennard-Jones interactions were gradually decoupled, and because the azobenzene molecule carries no charge, electrostatic interactions were not scaled. Simulations were minimized for 500 steps and equilibrated for 10 ns. Each $\lambda$ point was then run for 40 ns using a stochastic integrator. A soft-core potential ($\alpha = 0.5$, power $= 1$) was employed to avoid singularities when interactions were switched off.

The free energies were calculated using the Python package Alchemlyb[2] and the multi-state Bennett acceptance ratio.[3] The octanol-water partitioning logP was calculated according to the following equation:

$$logP = \frac{\Delta G_{W}^{solv} - \Delta G_{OCO}^{solv}}{RT \cdot ln(10)}$$

### 2 Azobenzene Parametrization

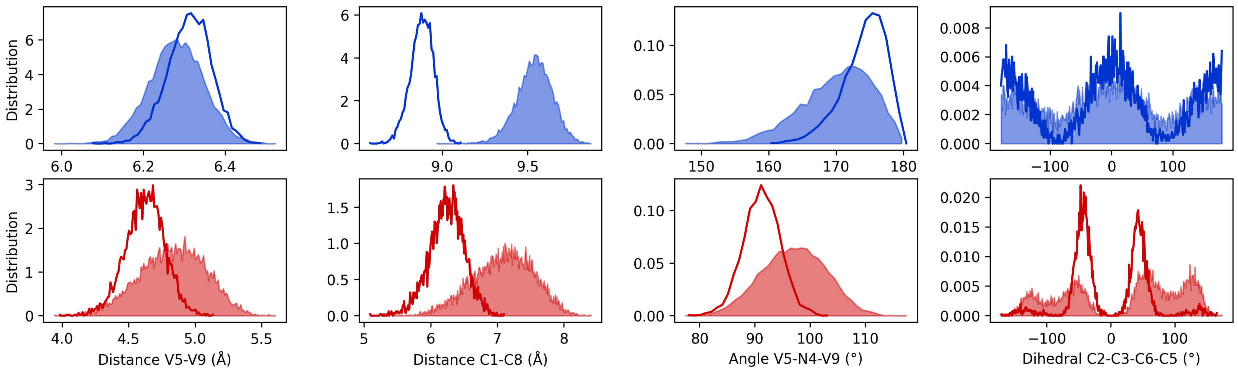


Figure S1: Comparison of Bonded Distributions between semi-empirical QM calculations using Bartender (filled) and classical MD Simulations using CHARMM36 (lines) pseudo-CG (mapped from AA) trajectories. The distributions of the trans (blue) and cis (red) isomer of azobenzene are shown.

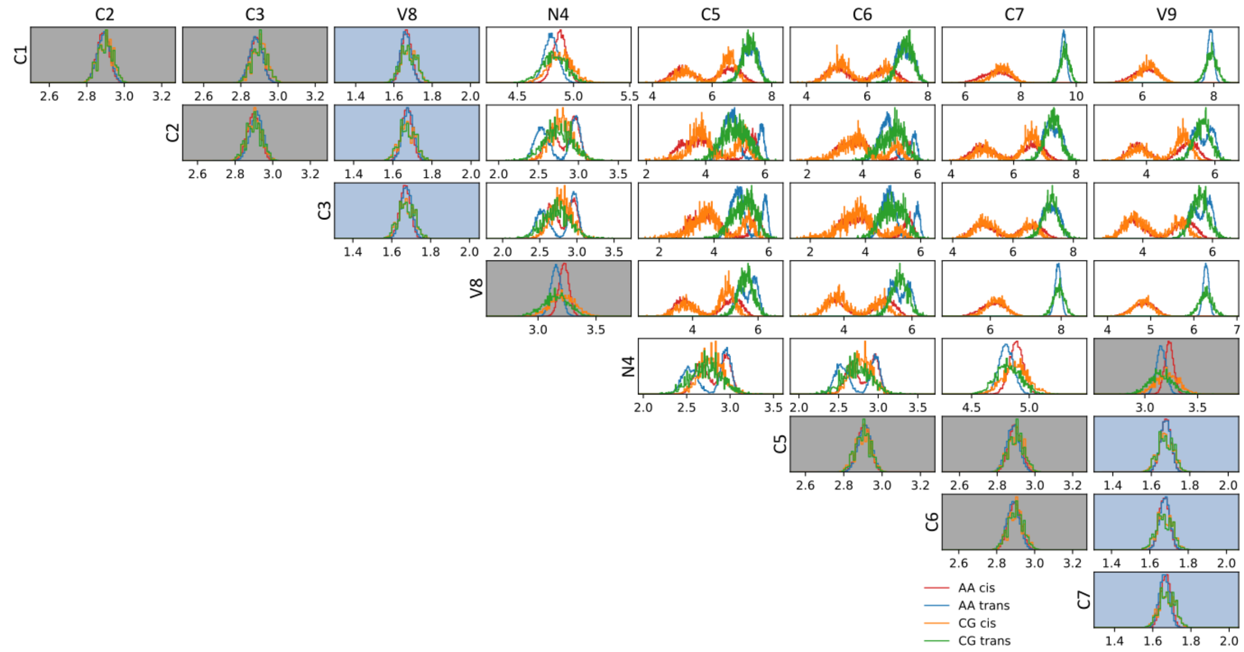


Figure S2: Distance distributions of the CG Martini 3 model for azobenzene. Distributions of all distances from pseudo-CG (mapped from AA) and CG trajectories are shown. Distances are in Ångström. Highlighted are interactions defined using bonds or constraints (grey) and interactions involved in the construction of virtual sites (blue).

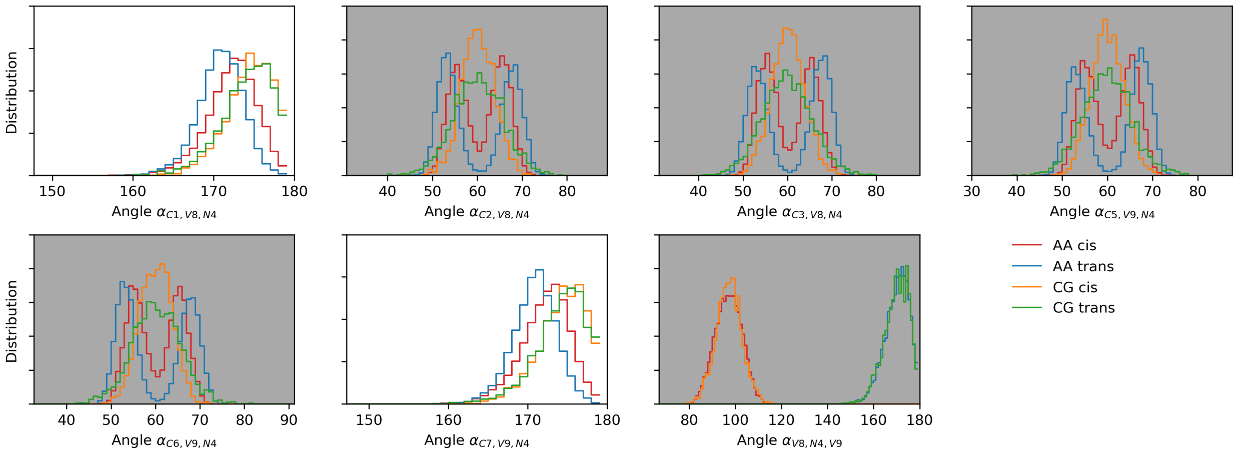


Figure S3: Angle distributions of the CG Martini 3 model for azobenzene. Distributions of angles from pseudo-CG (mapped from AA) and CG trajectories are shown. Highlighted are interactions defined using angle potentials (grey).

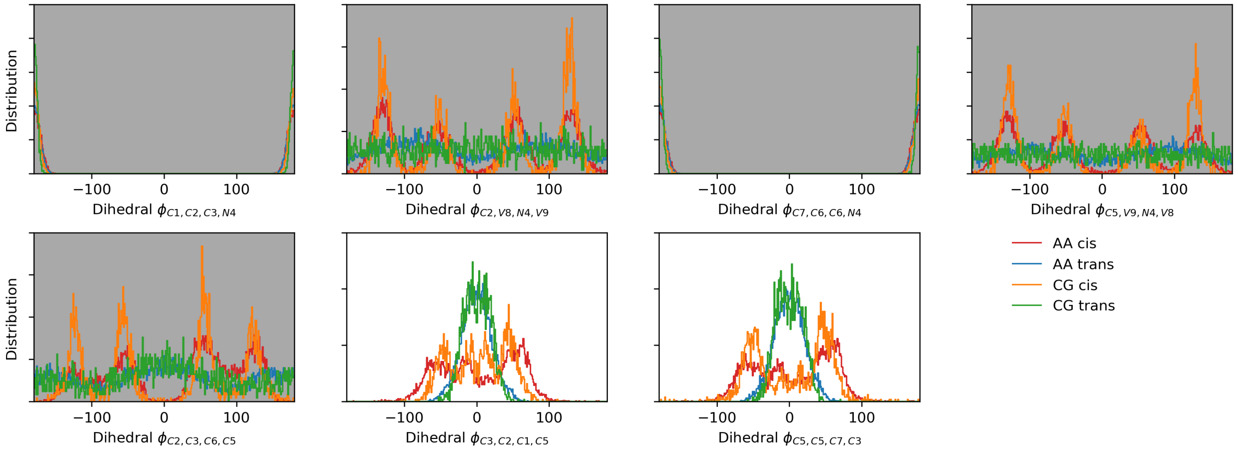


Figure S4: Dihedral distributions of the CG Martini 3 model for azobenzene. Distributions of dihedrals from pseudo-CG (mapped from AA) and CG trajectories are shown. Highlighted are interactions defined using dihedral potentials (grey).

### 3 Bonded terms

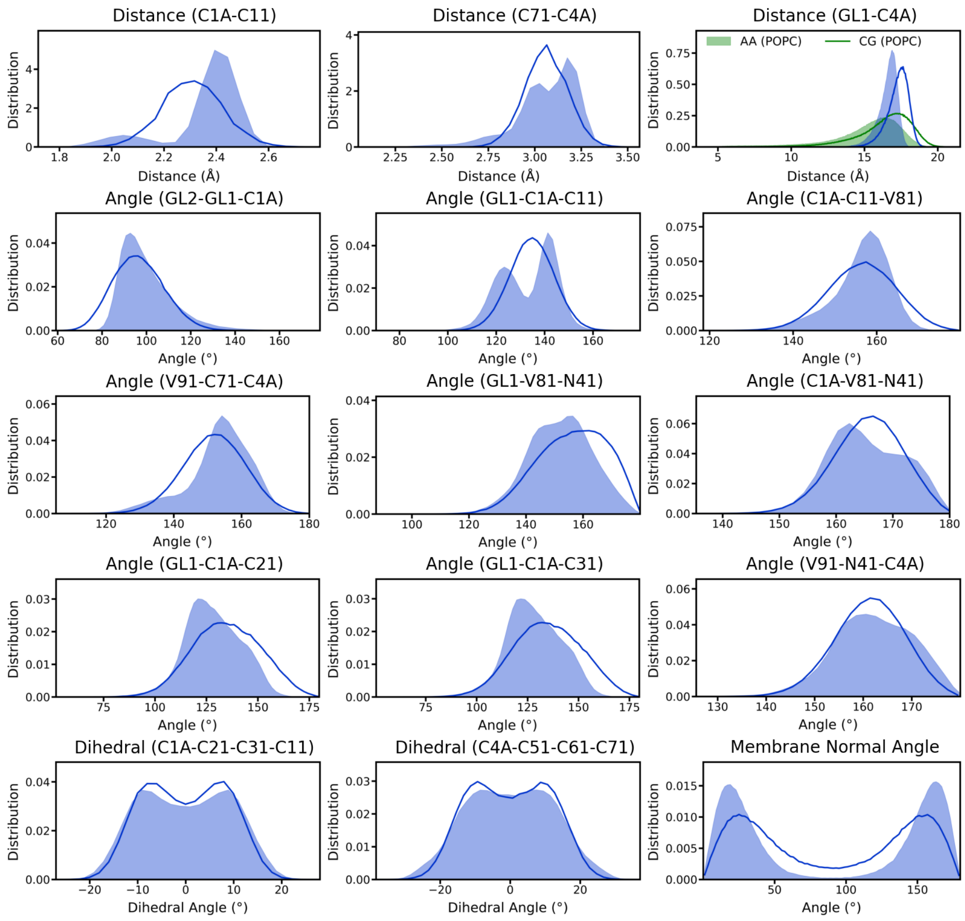


Figure S5: Bonded distributions of the CG Martini 3 model for STAPC. Distributions of selected distances, angles, and dihedral angles from pseudo-CG (mapped from AA, filled) and CG trajectories (lines) are shown including all modified bonded terms introduced or adapted to capture the geometry and conformational rigidity of the azobenzene-containing lipid tail (i.e., bonds C1A–C11 and C71–C4A, angles GL1-C1A-C11 and V91-C71-C4A, and dihedral angle C1A-C21-C31-C11). Further distributions are shown for validation. The GL1–C4A distance distributions (top right) illustrate the effective length of the photoswitchable tail in comparison to POPC (green) as reference. Angle distributions for the orientation of azobenzene with respect to the membrane normal are shown on the bottom right.

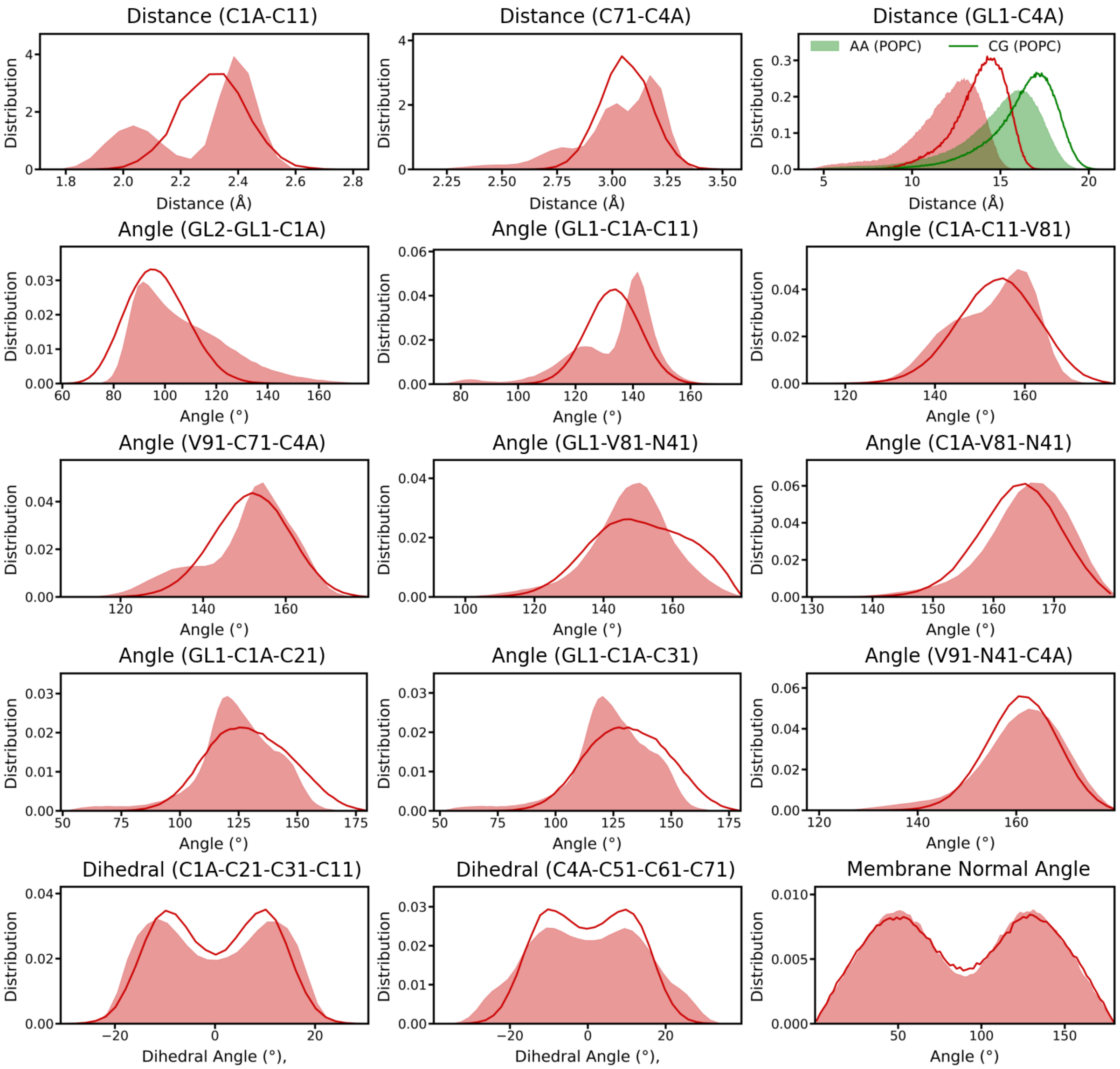


Figure S6: Bonded distributions of the CG Martini 3 model for SCAPC. Distributions of selected distances, angles, and dihedral angles from pseudo-CG (mapped from AA, filled) and CG trajectories (lines) are shown including all modified bonded terms introduced or adapted to capture the geometry and conformational rigidity of the azobenzene-containing lipid tail (i.e., bonds C1A–C11 and C71–C4A, angles GL1-C1A-C11 and V91-C71-C4A, and dihedral angle C1A-C21-C31-C11). Further distributions are shown for validation. The GL1–C4A distance distributions (top right) illustrate the effective length of the photoswitchable tail in comparison to POPC (green) as reference. Angle distributions for the orientation of azobenzene with respect to the membrane normal are shown on the bottom right.

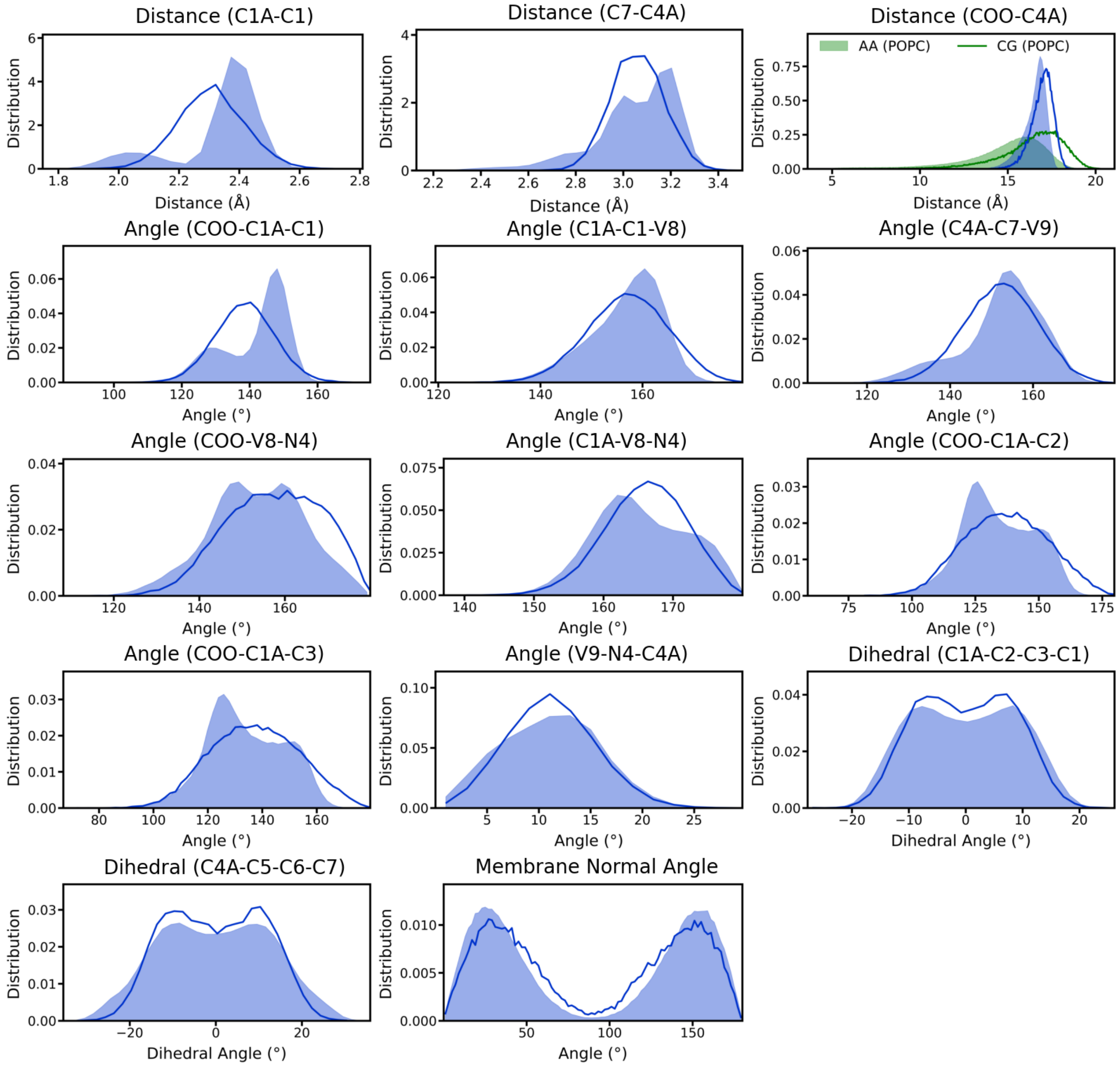


Figure S7: Bonded distributions of the CG Martini 3 model for TAFA. Distributions of selected distances, angles, and dihedral angles from pseudo-CG (mapped from AA, filled) and CG trajectories (lines) are shown including all modified bonded terms introduced or adapted to capture the geometry and conformational rigidity of the azobenzene-containing lipid tail (i.e., bonds C1A–C1 and C7–C4A, angles GL1-C1A-C1 and V9-C7-C4A, and dihedral angle C1A-C2-C3-C1). Further distributions are shown for validation. The COO–C4A distance distributions (top right) illustrate the effective length of the photoswitchable tail in comparison to GL1-C4A from POPC (green) as reference. Angle distributions for the orientation of azobenzene with respect to the membrane normal are shown on the bottom right.

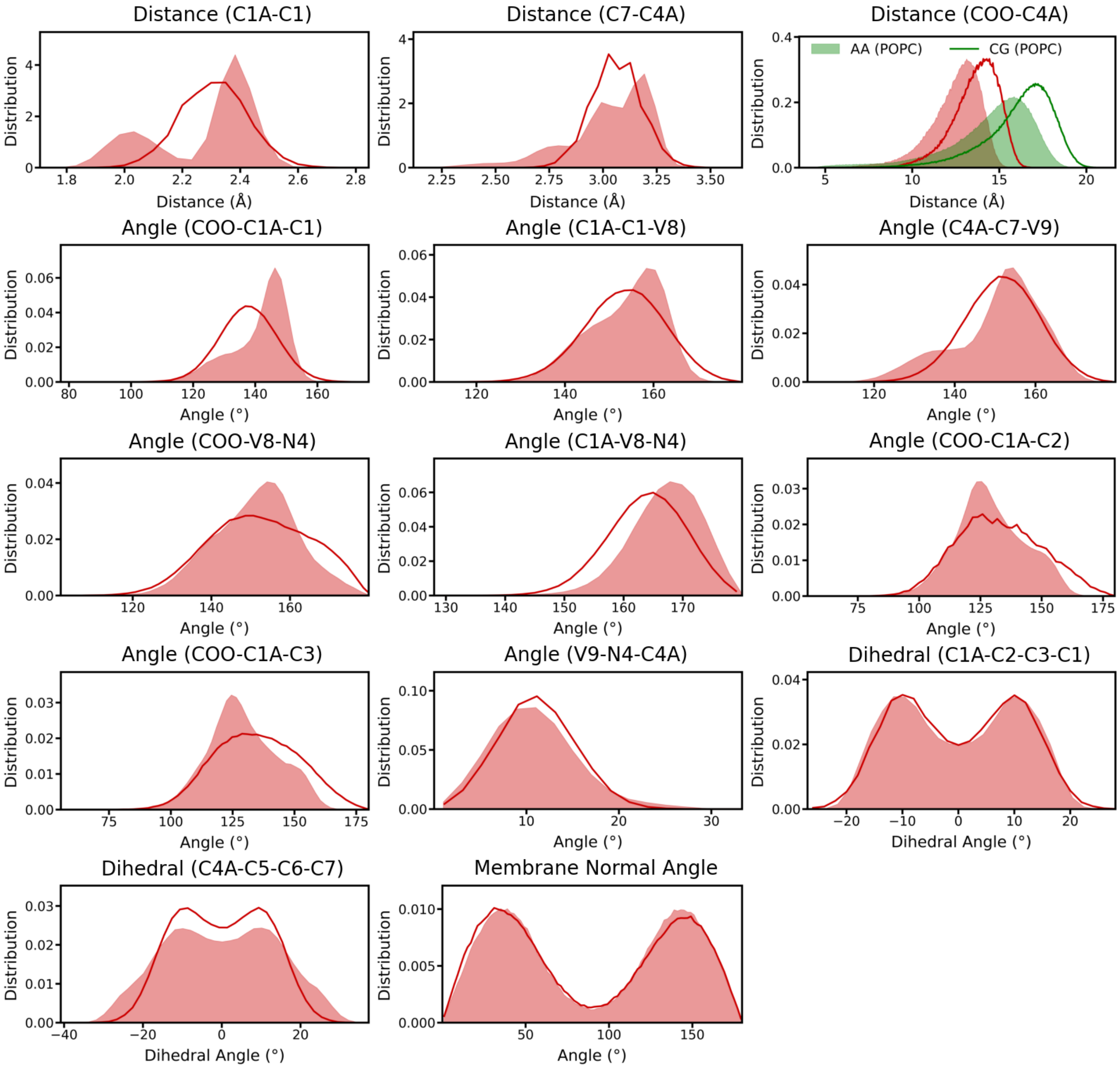


Figure S8: Bonded distributions of the CG Martini 3 model for CAFA. Distributions of selected distances, angles, and dihedral angles from pseudo-CG (mapped from AA, filled) and CG trajectories (lines) are shown including all modified bonded terms introduced or adapted to capture the geometry and conformational rigidity of the azobenzene-containing lipid tail (i.e., bonds C1A–C1 and C7–C4A, angles GL1-C1A-C1 and V9-C7-C4A, and dihedral angle C1A-C2-C3-C1). Further distributions are shown for validation. The COO–C4A distance distributions (top right) illustrate the effective length of the photoswitchable tail in comparison to GL1-C4A from POPC (green) as reference. Angle distributions for the orientation of azobenzene with respect to the membrane normal are shown on the bottom right.

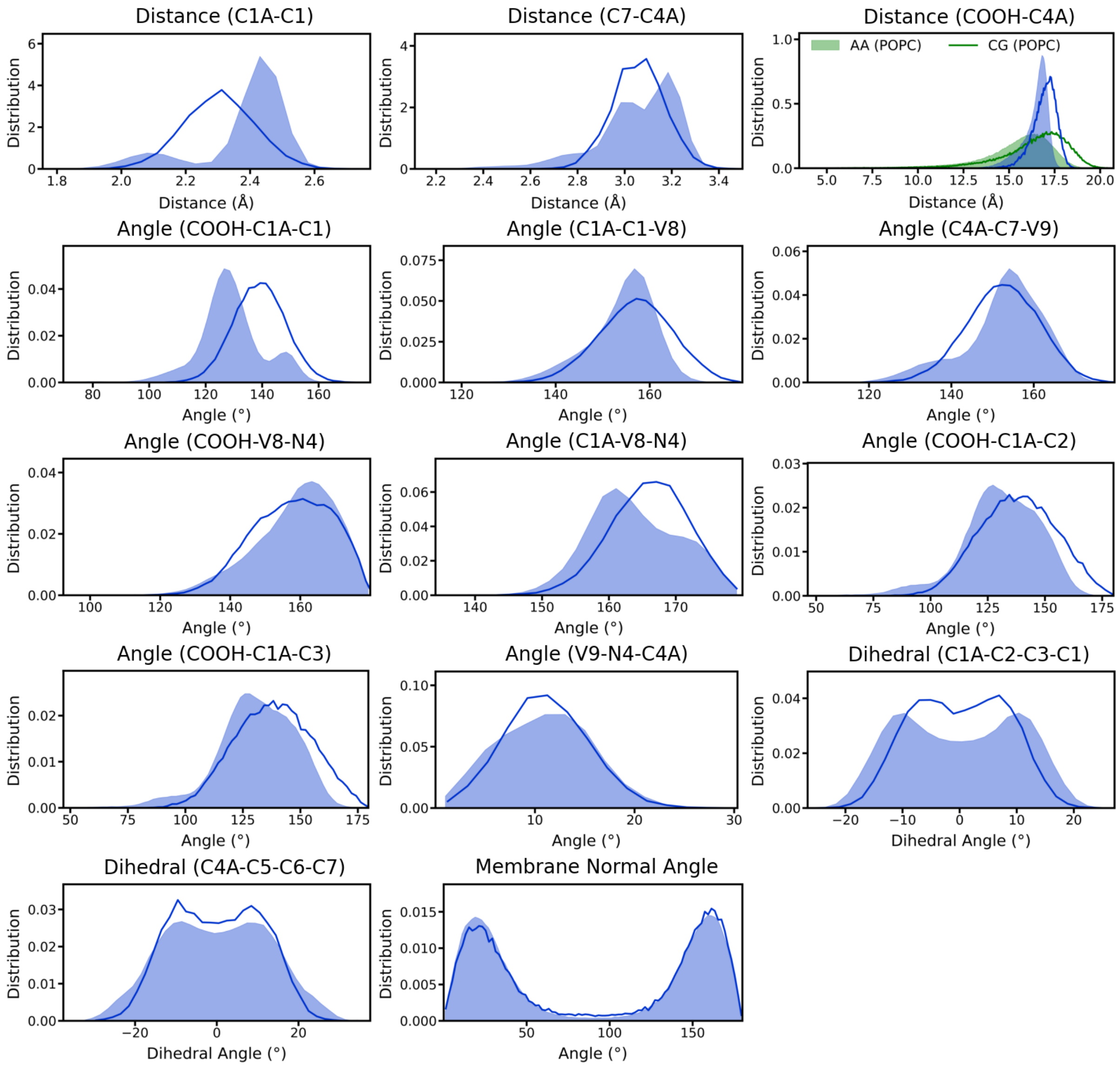


Figure S9: Bonded distributions of the CG Martini 3 model for TAFAH. Distributions of selected distances, angles, and dihedral angles from pseudo-CG (mapped from AA, filled) and CG trajectories (lines) are shown including all modified bonded terms introduced or adapted to capture the geometry and conformational rigidity of the azobenzene-containing lipid tail (i.e., bonds C1A–C1 and C7–C4A, angles GL1-C1A-C1 and V9-C7-C4A, and dihedral angle C1A-C2-C3-C1). Further distributions are shown for validation. The COOH–C4A distance distributions (top right) illustrate the effective length of the photoswitchable tail in comparison to GL1-C4A from POPC (green) as reference. Angle distributions for the orientation of azobenzene with respect to the membrane normal are shown on the bottom right.

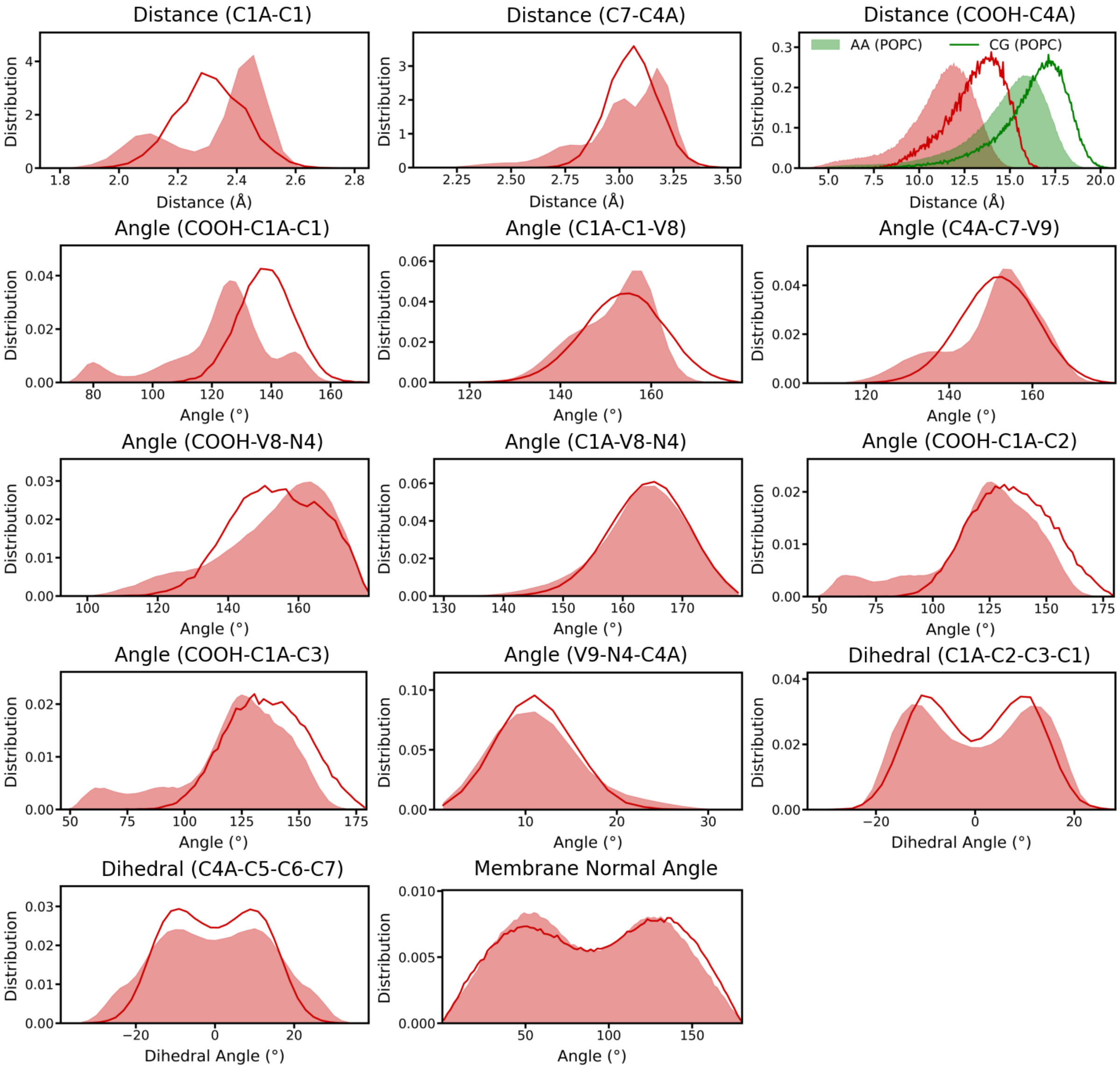


Figure S10: Bonded distributions of the CG Martini 3 model for CAFAH. Distributions of selected distances, angles, and dihedral angles from pseudo-CG (mapped from AA, filled) and CG trajectories (lines) are shown including all modified bonded terms introduced or adapted to capture the geometry and conformational rigidity of the azobenzene-containing lipid tail (i.e., bonds C1A–C1 and C7–C4A, angles GL1-C1A-C1 and V9-C7-C4A, and dihedral angle C1A-C2-C3-C1). Further distributions are shown for validation. The COOH–C4A distance distributions (top right) illustrate the effective length of the photoswitchable tail in comparison to GL1-C4A from POPC (green) as reference. Angle distributions for the orientation of azobenzene with respect to the membrane normal are shown on the bottom right.

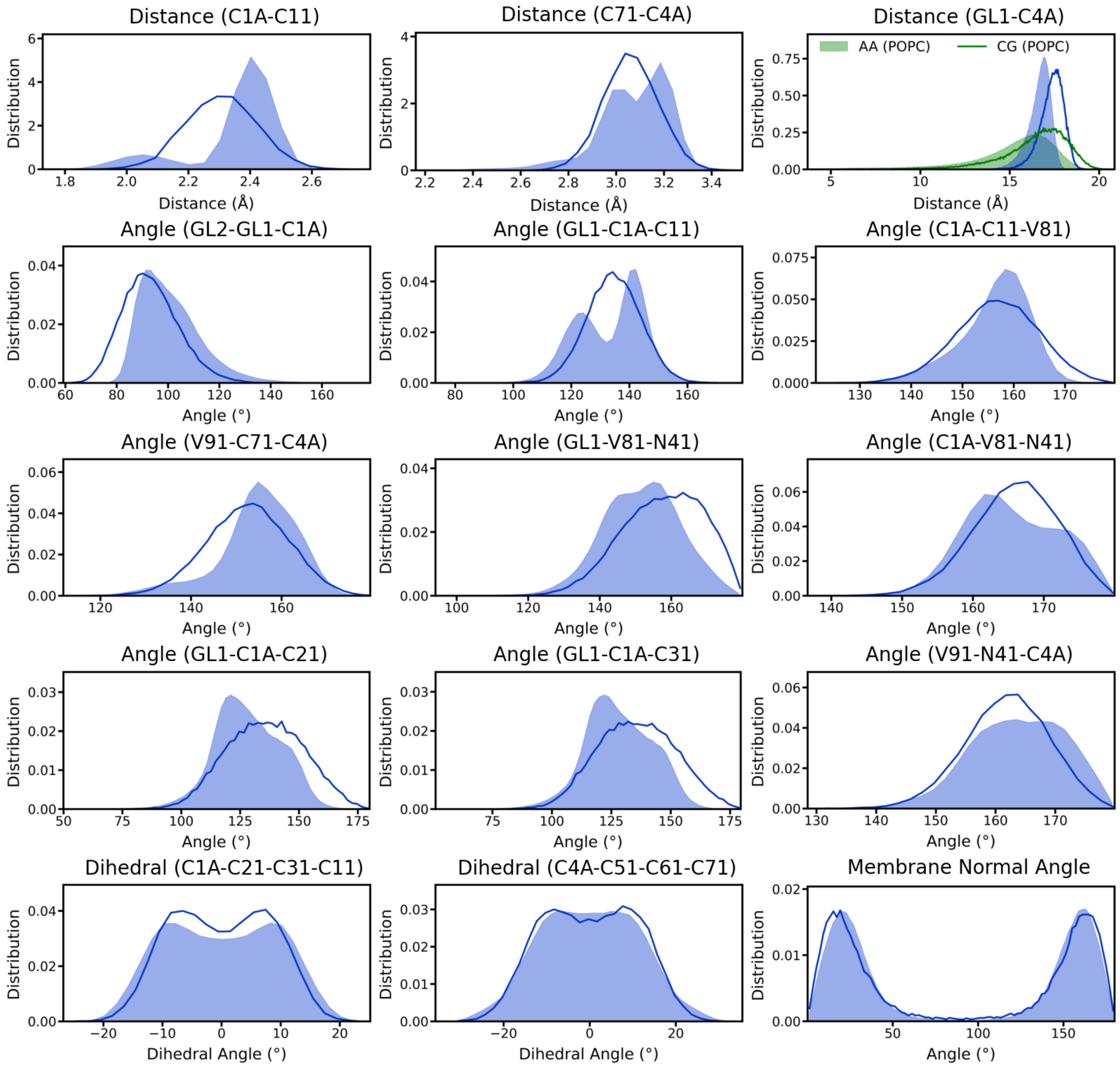


Figure S11: Bonded distributions of the CG Martini 3 model for the first tail DTAPC. Distributions of selected distances, angles, and dihedral angles from pseudo-CG (mapped from AA, filled) and CG trajectories (lines) are shown including all modified bonded terms introduced or adapted to capture the geometry and conformational rigidity of the first azobenzene-containing lipid tail (i.e., bonds C1A–C11 and C71–C4A, angles GL1-C1A-C11 and V91-C71-C4A, and dihedral angle C1A-C21-C31-C11). Further distributions are shown for validation. The GL1–C4A distance distributions (top right) illustrate the effective length of the first photoswitchable tail in comparison to POPC (green) as reference. Angle distributions for the orientation of azobenzene with respect to the membrane normal are shown on the bottom right.

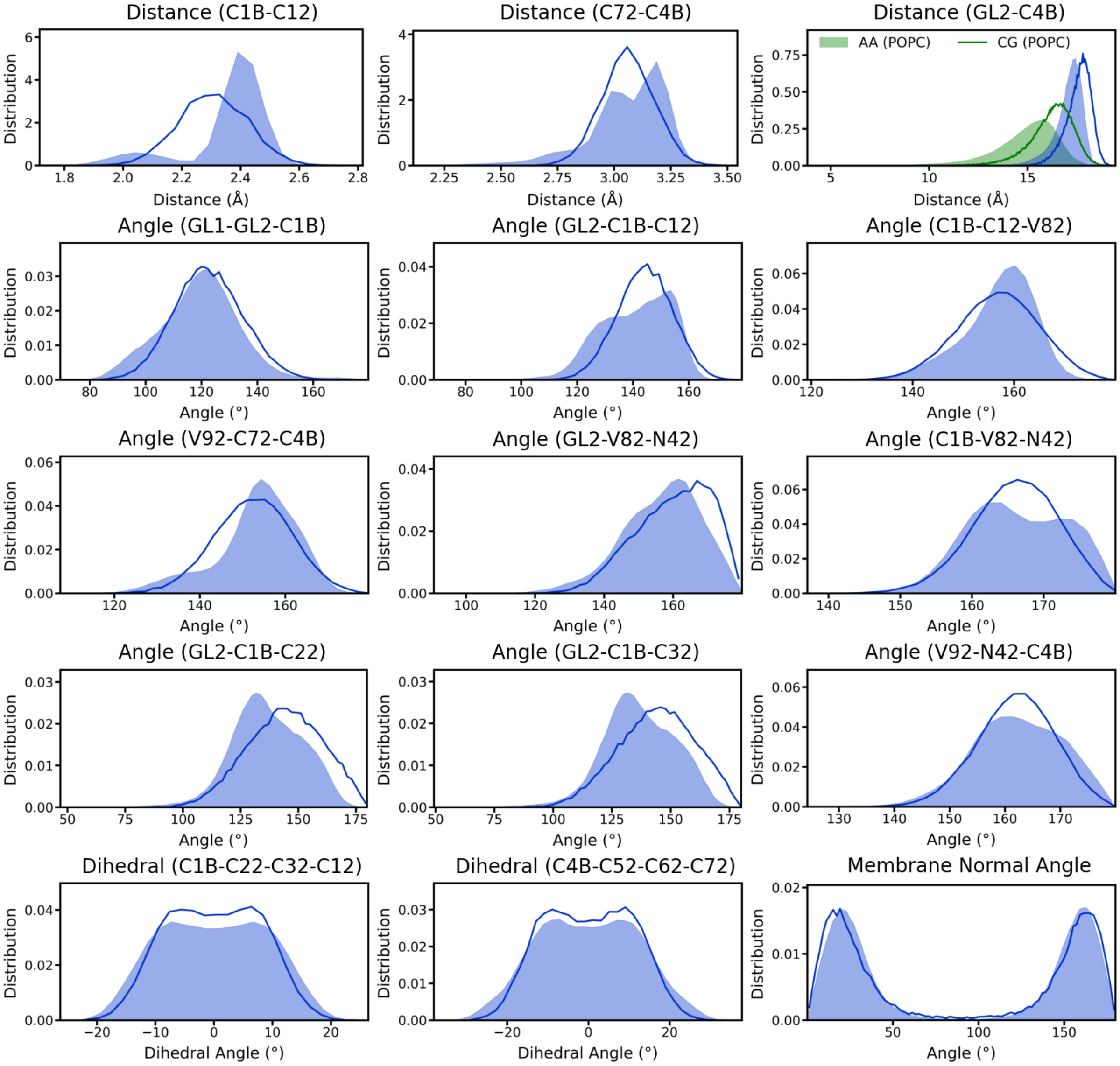


Figure S12: Bonded distributions of the CG Martini 3 model for the second tail DTAPC. Distributions of selected distances, angles, and dihedral angles from pseudo-CG (mapped from AA, filled) and CG trajectories (lines) are shown including all modified bonded terms introduced or adapted to capture the geometry and conformational rigidity of the second azobenzene-containing lipid tail (i.e., bonds C1B–C12 and C72–C4B, angles GL2-C1B-C12 and V92-C72-C4B, and dihedral angle C1B-C22-C32-C12). Further distributions are shown for validation. The GL2–C4B distance distributions (top right) illustrate the effective length of the second photoswitchable tail in comparison to POPC (green) as reference. Angle distributions for the orientation of azobenzene with respect to the membrane normal are shown on the bottom right.

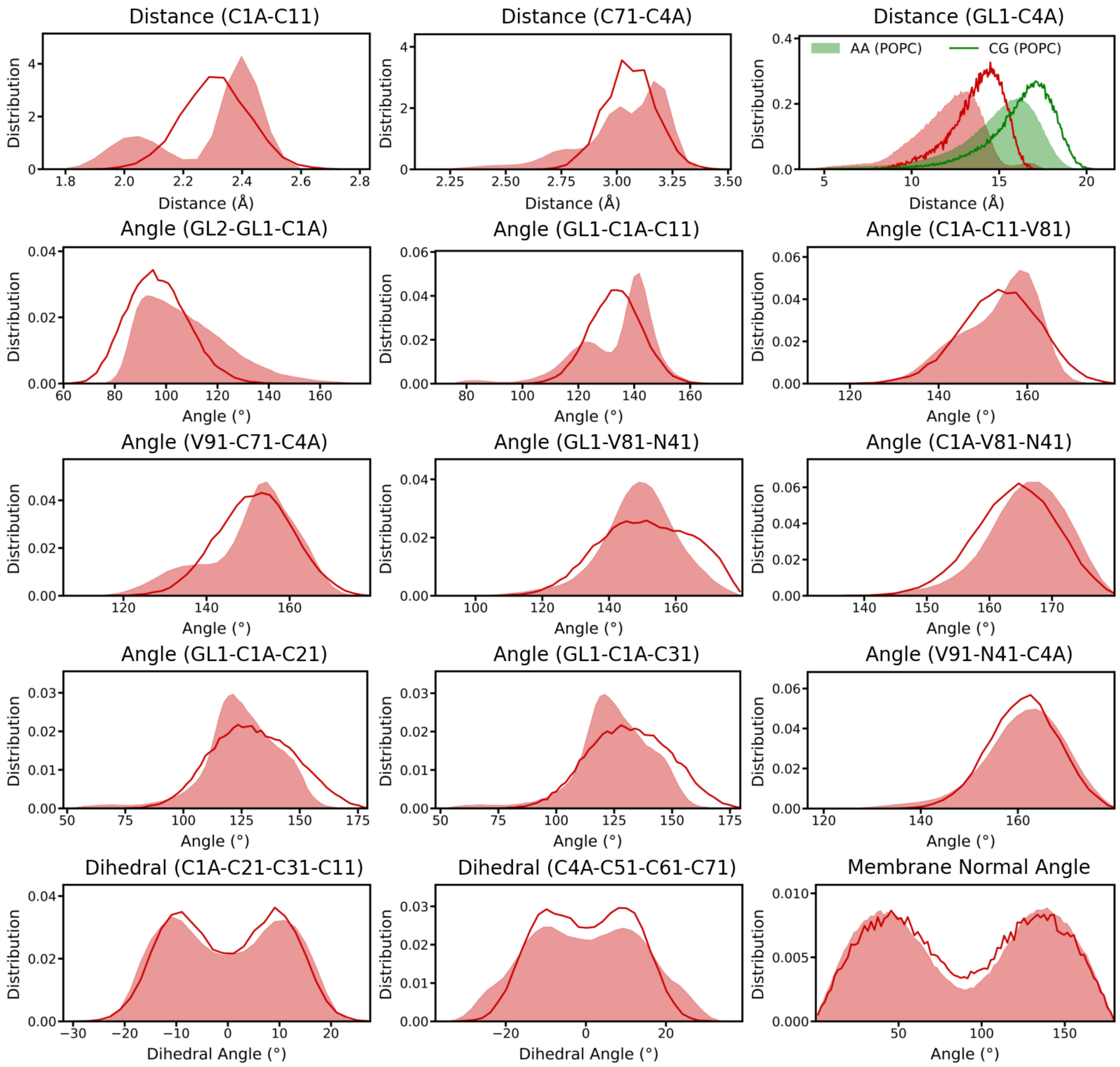


Figure S13: Bonded distributions of the CG Martini 3 model for the first tail DCAPC. Distributions of selected distances, angles, and dihedral angles from pseudo-CG (mapped from AA, filled) and CG trajectories (lines) are shown including all modified bonded terms introduced or adapted to capture the geometry and conformational rigidity of the first azobenzene-containing lipid tail (i.e., bonds C1A–C11 and C71–C4A, angles GL1-C1A-C11 and V91-C71-C4A, and dihedral angle C1A-C21-C31-C11). Further distributions are shown for validation. The GL1–C4A distance distributions (top right) illustrate the effective length of the first photoswitchable tail in comparison to POPC (green) as reference. Angle distributions for the orientation of azobenzene with respect to the membrane normal are shown on the bottom right.

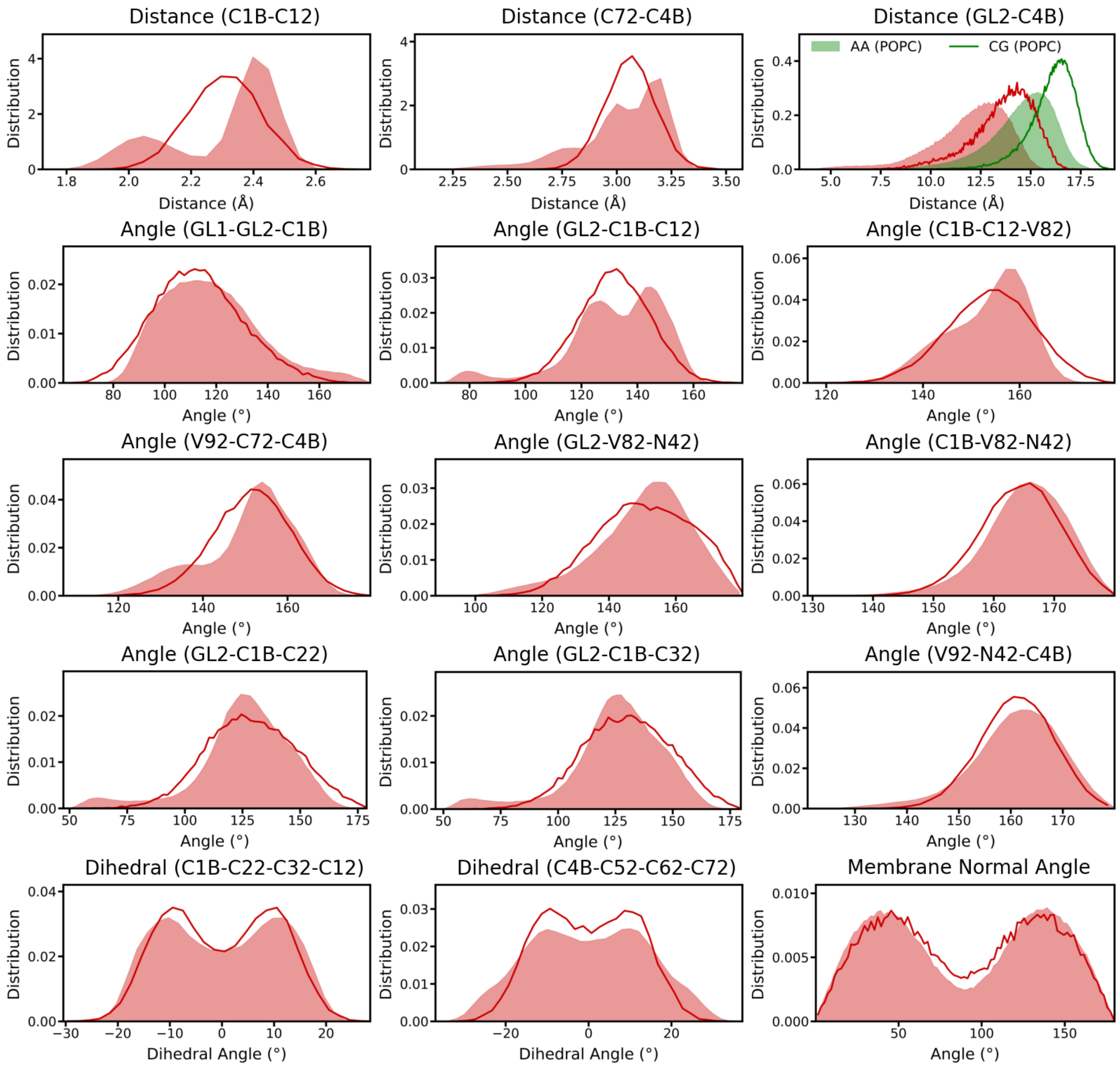


Figure S14: Bonded distributions of the CG Martini 3 model for the second tail DCAPC. Distributions of selected distances, angles, and dihedral angles from pseudo-CG (mapped from AA, filled) and CG trajectories (lines) are shown including all modified bonded terms introduced or adapted to capture the geometry and conformational rigidity of the second azobenzene-containing lipid tail (i.e., bonds C1B–C12 and C72–C4B, angles GL2-C1B-C12 and V92-C72-C4B, and dihedral angle C1B-C22-C32-C12). Further distributions are shown for validation. The GL2–C4B distance distributions (top right) illustrate the effective length of the second photoswitchable tail in comparison to POPC (green) as reference. Angle distributions for the orientation of azobenzene with respect to the membrane normal are shown on the bottom right.

### 4 Density plots

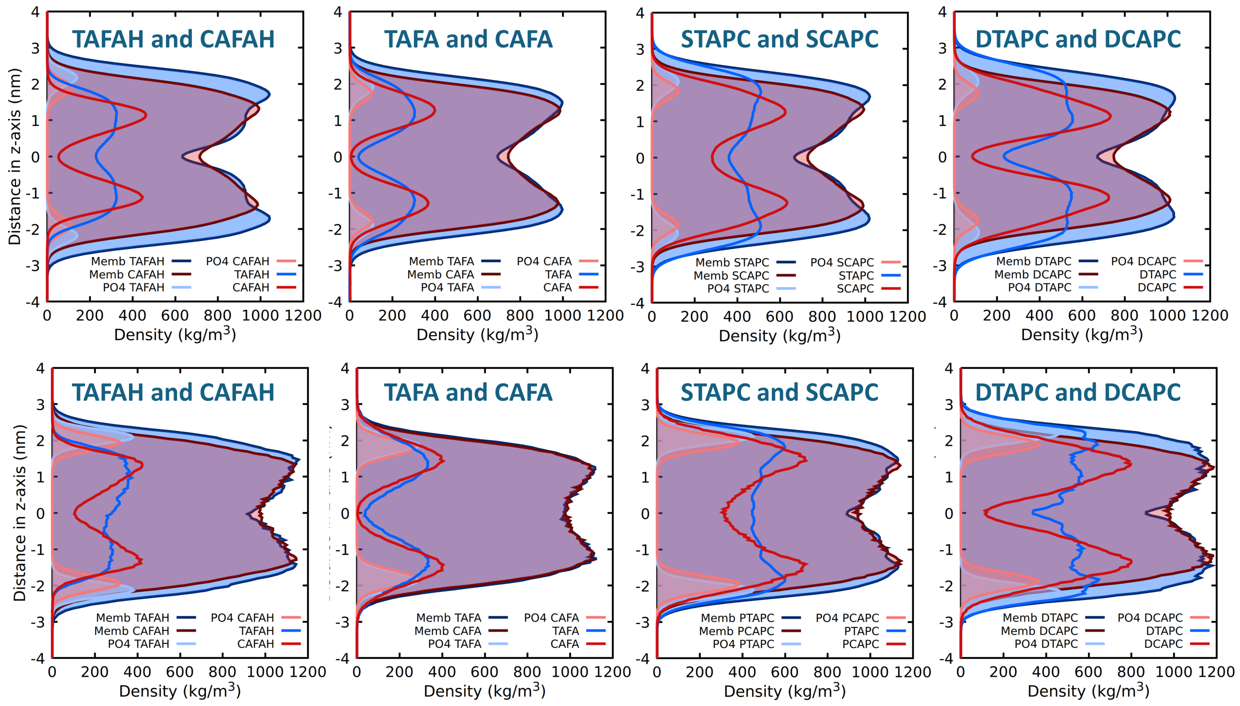


Figure S15: Mass densities of the whole membrane, phosphate group, and the photolipids (T/C)AFA, (T/C)AFAH, S(T/C)APC and D(T/C)APC (photolipids scaled by a factor of two) at 303K for the AA (upper panel) and CG simulation (lower panel) along the z axis.

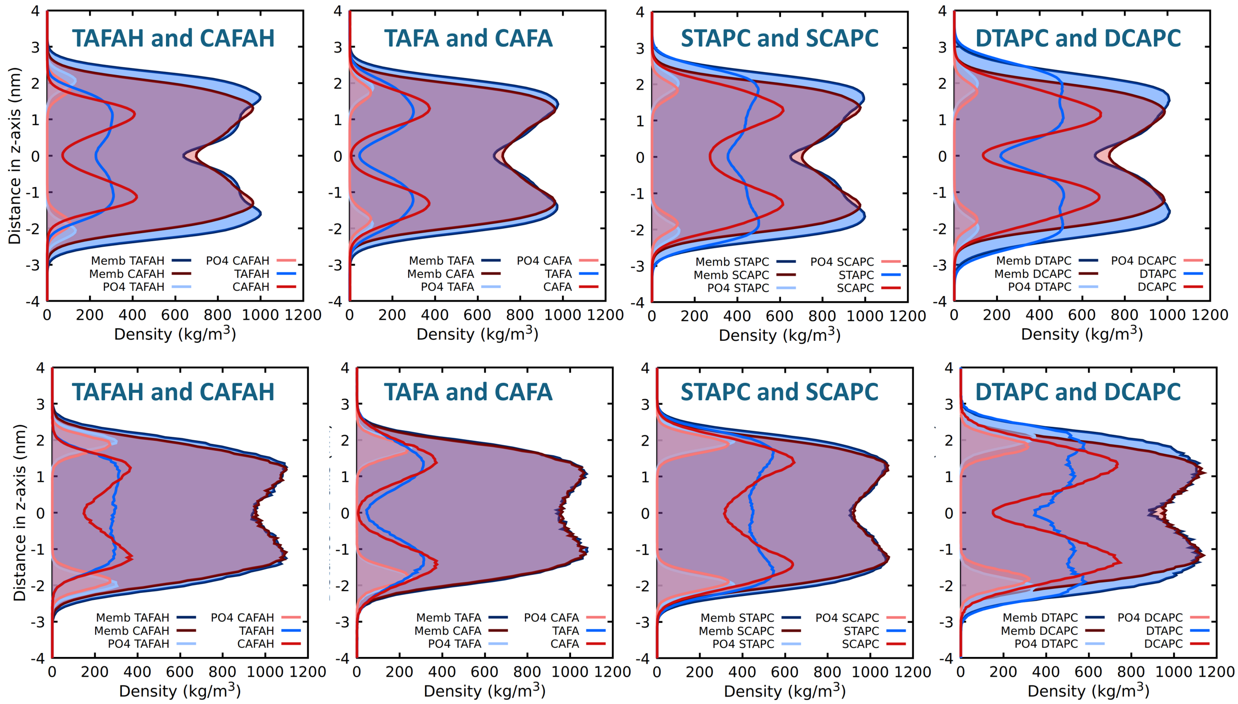


Figure S16: Mass densities of the whole membrane, phosphate group, and the photolipids (T/C)AFA, (T/C)AFAH, S(T/C)APC and D(T/C)APC (photolipids scaled by a factor of two) at 330K for the AA (upper panel) and CG simulation (lower panel) along the z axis.

### 5 Membrane thickness and area per lipid analysis

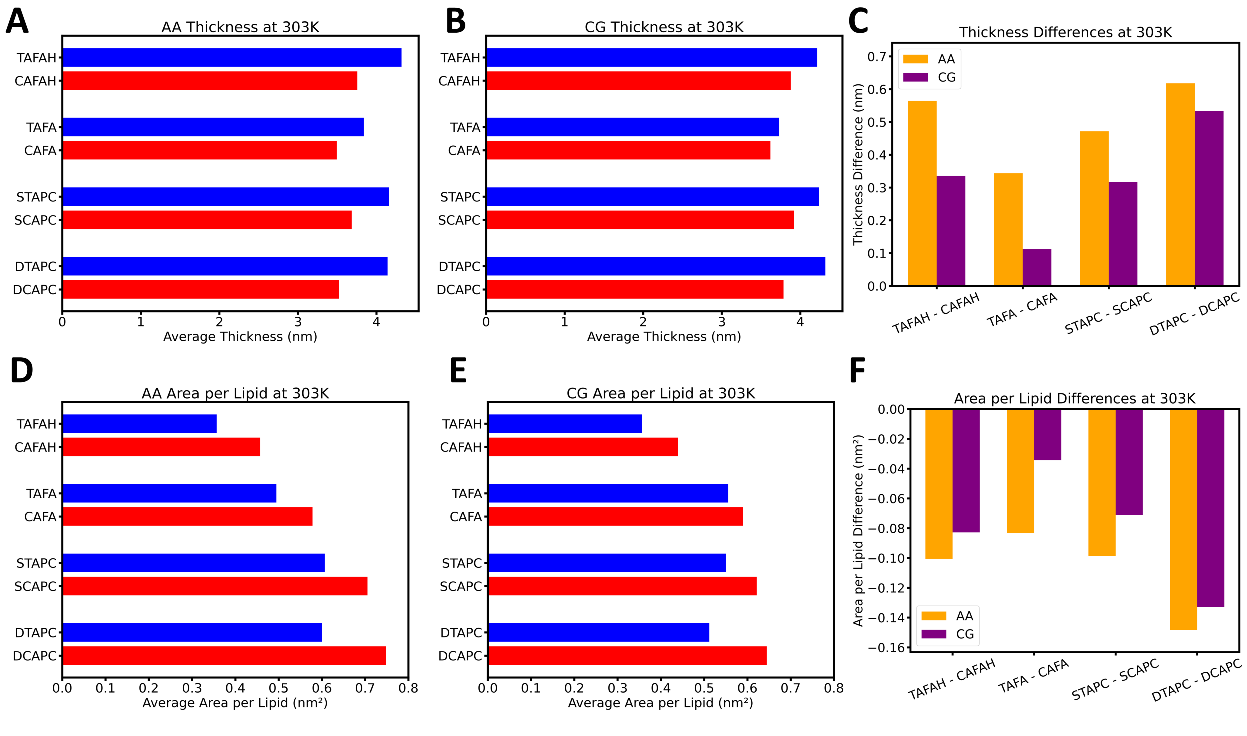


Figure S17: Properties for binary lipid mixtures containing 25 mol% azobenzene-based photolipid and 75 mol% POPC at 303K. (A,B) Membrane thickness measured as the average z distance of the P atoms (AA, A) and of the PO4 beads (CG, B), respectively. The photolipids are protonated (CAFAH/TAFAH) and deprotonated fatty acid (CAFA/TAFA), PC with one photoswitchable tail (SCAPC/STAPC), and PC with two photoswitchable tails (DCAPC/DTAPC). (C) Thickness difference between *trans*- and *cis*-isomer of each photolipid species for AA (orange) and CG systems (purple). (D,E) Area per lipid of the photolipids for the AA (D) and CG systems (E). (F) Area per lipid difference between *trans*- and *cis*-isomer of each photolipid species for AA (orange) and CG systems (purple).

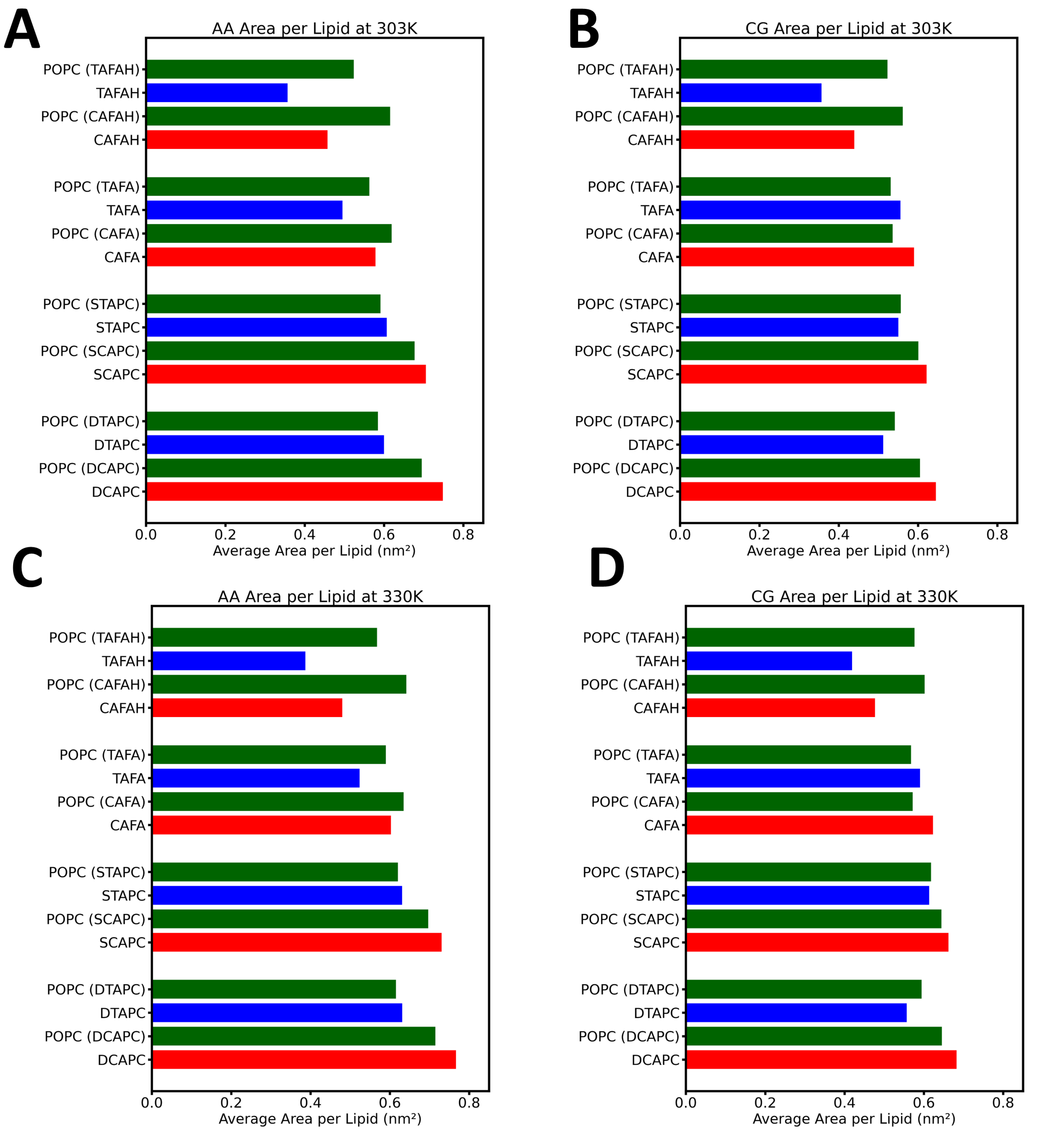


Figure S18: Area per lipid of the photolipids and POPC of each system at 303K for the AA (A) and CG (B), and at 330K for the AA (C) and CG (D).The photolipids are protonated (CAFAH/TAFAH) and deprotonated fatty acid (CAFA/TAFA), PC with one photoswitchable tail (SCAPC/STAPC), and PC with two photoswitchable tails (DCAPC/DTAPC).

#### 6 Comparative snapshots of (T/C)AFAH and S(T/C)APC

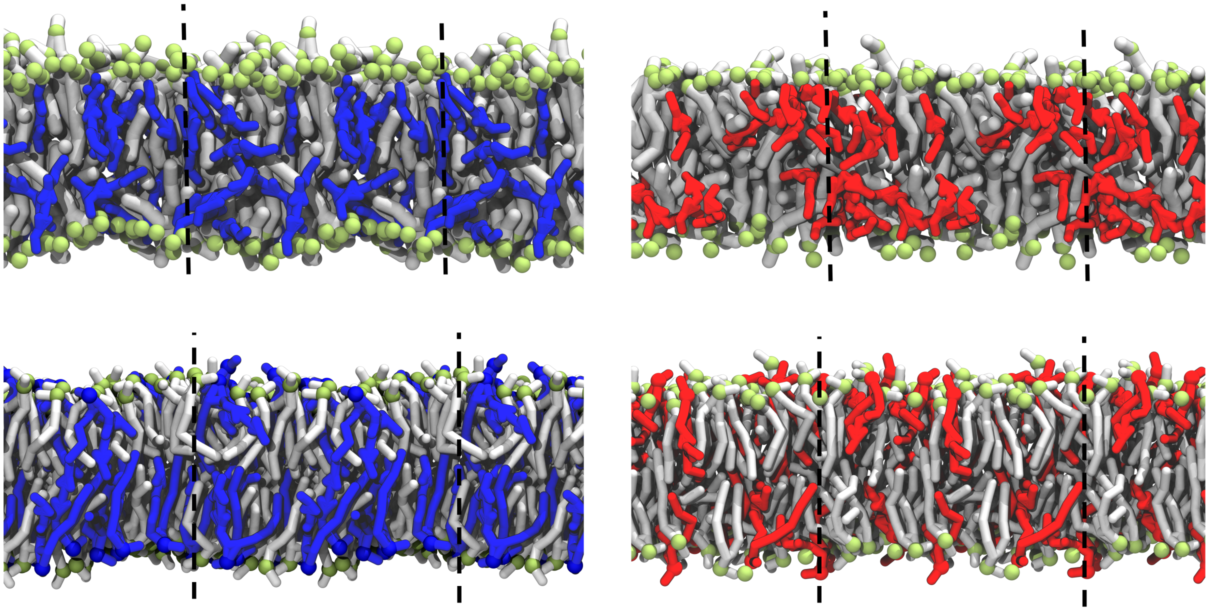

Figure S19: Snapshots from the simulations of TAFAH (top left), CAFAH (top right), STAPC (bottom left), and SCAPC (bottom right). The simulation box is indicated by dashed black lines. Periodic images of the system are shown to illustrate the continuity due to periodic boundary conditions.

### 7 DgkA flexibility of the three replicas

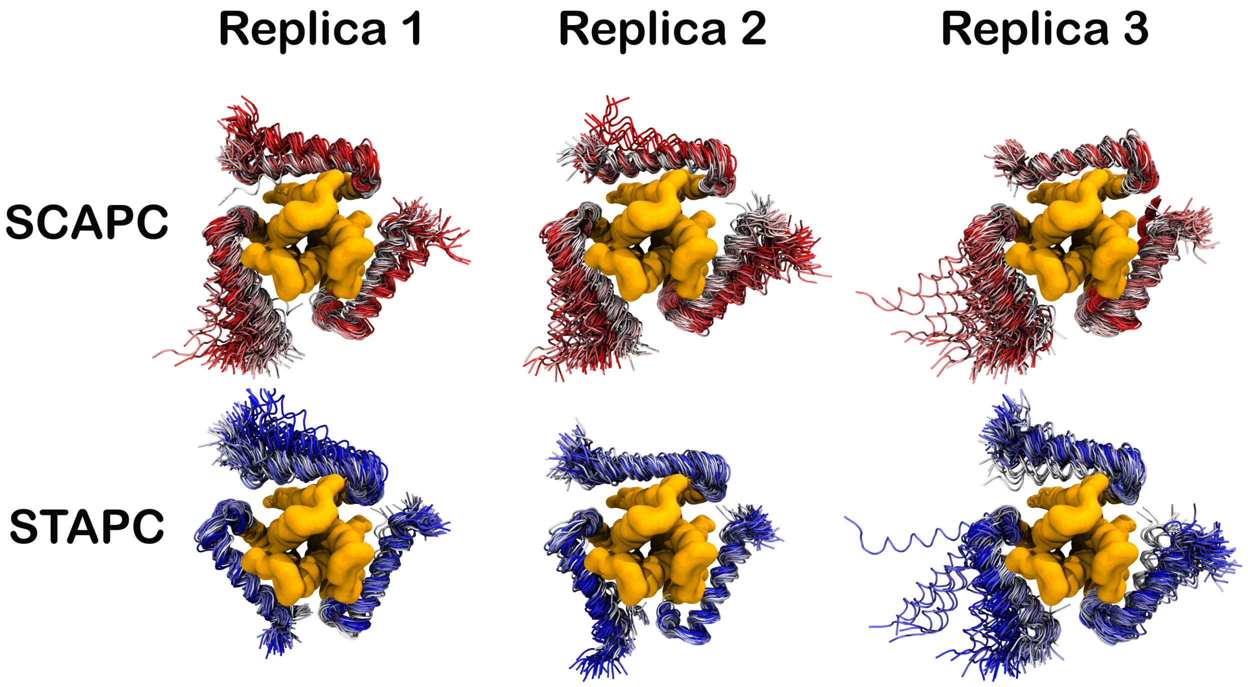


Figure S20: Movement of the surface helix of a DgkA trimer in the three replicas of a SCAPC- (top row) and STAPC-containing (bottom row) membrane. 100 equally spaced frames of a 10 µs trajectory are depicted.

### 8 US convergence tests and histograms

The convergence tests show the PMF computed for increasing chunks of 100 ns starting from 200 ns to 500 ns, considering the equilibration of the system to be finished after 200 ns.

The histograms show the bins of each window in different colours and the sum of them is displayed with a black line.

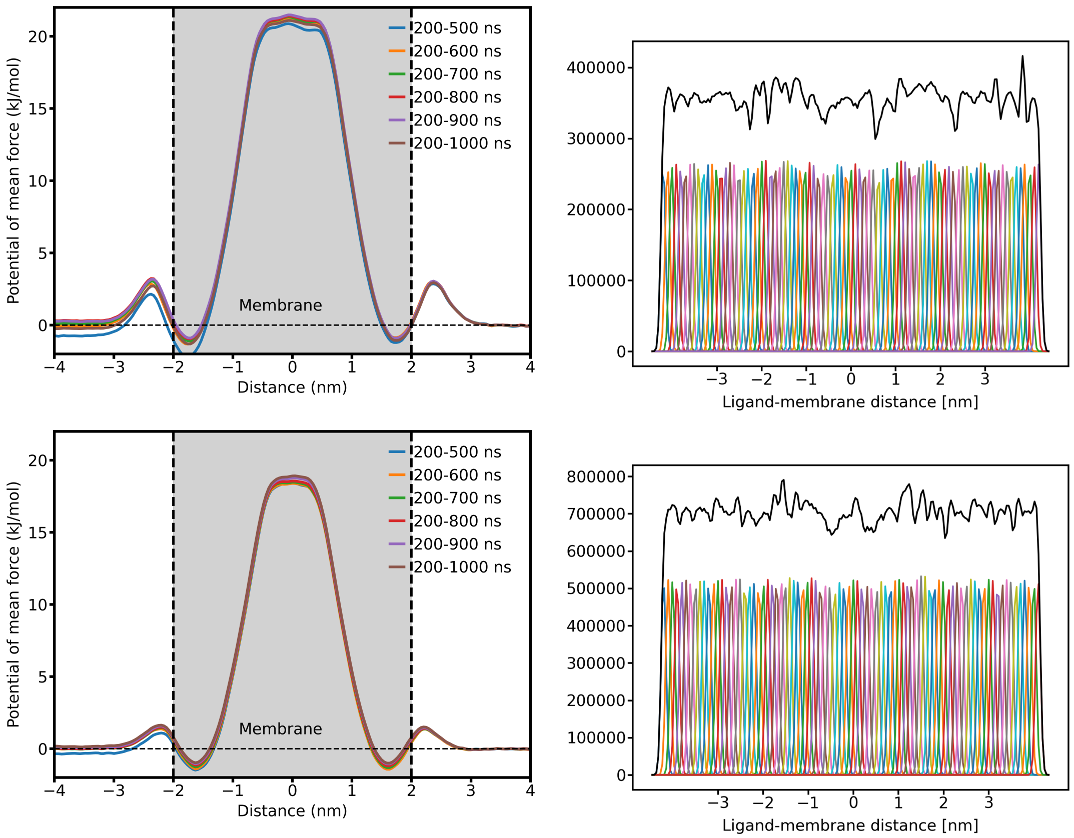


Figure S21: (Top, left) Convergence test and (top, right) histogram of the US of the salbutamol molecule pulled along the DTAPC-POPC membrane. (85 windows, every 0.1 nm, 1 µs per window). (Bottom, left) Convergence test and (bottom, right) histogram of the US of the salbutamol molecule pulled along the DCAPC-POPC membrane. (85 windows, every 0.1 nm, 1 µs per window).

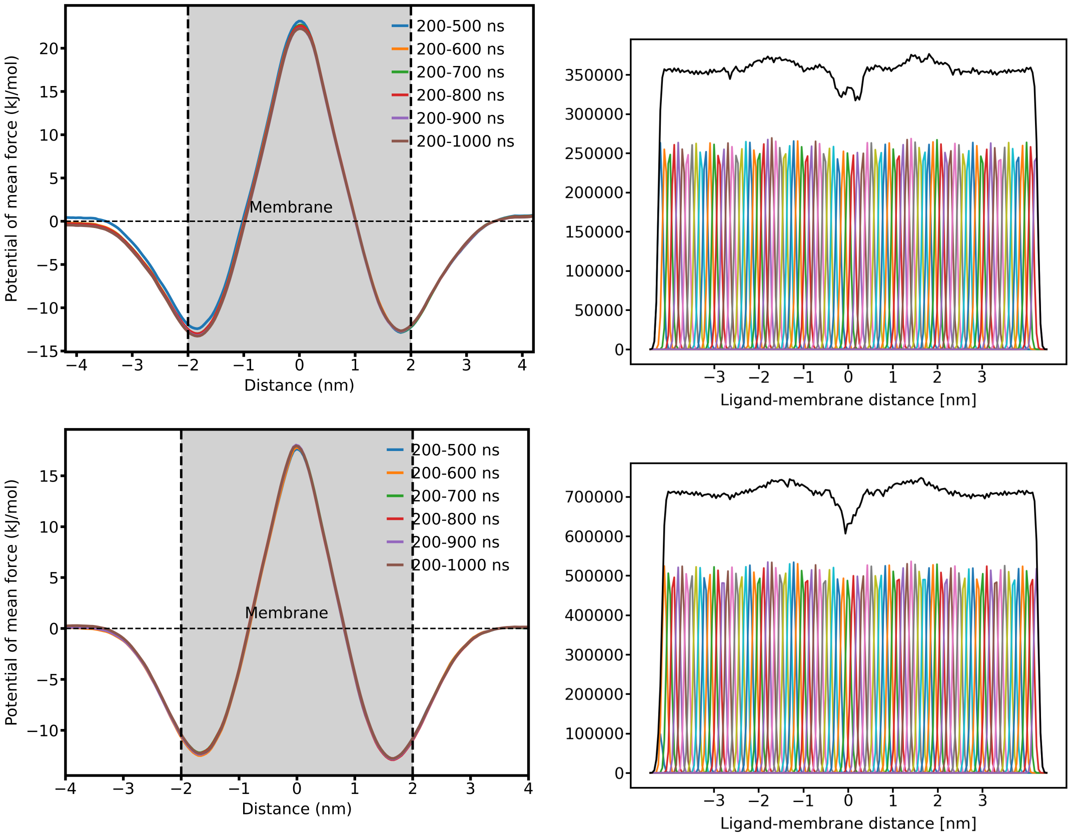


Figure S22: (Top, left) Convergence test and (top, right) histogram of the US of the baricitinib molecule pulled along the DTAPC-POPC membrane. (85 windows, every 0.1 nm, 1 µs per window). (Bottom, left) Convergence test and (bottom, right) histogram of the US of the baricitinib molecule pulled along the DCAPC-POPC membrane. (85 windows, every 0.1 nm, 1 µs per window).
